# Pollution and Anthropogenic Stressors Are Associate with Cetacean Vulnerability in Coastal Waters: Fine-Scale Diagnostics from eDNA and Multispecies Modeling

**DOI:** 10.64898/2026.04.16.719104

**Authors:** Thilina S. Nimalrathna, Isis Guibert, Zhewei Si, Karen K. L. Yeung, Tsz Ying Chan, Mathew Seymour

## Abstract

Indo-Pacific humpback dolphin (*Sousa chinensis*) and finless porpoise (*Neophocaena phocaenoides*) are increasingly threatened across their native range, yet the relative influence of multiple stressors in shaping their population dynamics remains unclear. Current conservation strategies for both species are limited by incomplete data and limited assessment of affecting factors. Here, we integrated eDNA metabarcoding with Joint Species Distribution Modeling (JSDM) to assess how environmental gradients, pollution, and trophic associations interactively influence cetacean distributions in Hong Kong waters. We show that degraded water quality and intensified human activity negatively associated with cetacean occurrence, with clear species-specific responses to different stressors. *S. chinensis* covaried most strongly with Secchi disc depth, and presence of vessels, while *N. phocaenoides* was negatively associated with nitrate nitrogen and microbial pollution (sewage). The JSDM variance partitioning analysis highlighted that the occurrence of *S. chinensis* was primarily associated with anthropogenic disturbances (30.04%), followed by water physical properties (26.63%), whereas *N. phocaenoides* was more strongly associated with physical (40.9%) and anthropogenic disturbances (35.2%). By integrating eDNA and JSDM, our approach provides fine-scale diagnostics of species-specific vulnerabilities, supporting adaptive conservation strategies and guiding the realignment of protected areas to mitigate biodiversity loss in urbanized marine ecosystems.

**Environmental Implication:** Our study demonstrates that hazardous water pollutants, including microbial contamination, nutrient enrichment, and chemical stressors, vessel pressure, show strong, species-specific impacts on resident cetaceans in Hong Kong. By integrating eDNA metabarcoding with joint species distribution models, we provide a diagnostic framework that directly links pollutant profiles to ecological risk. These findings highlight that conventional conservation strategies overlooking pollution drivers are insufficient for marine megafauna persistence. Our approach offers an early-warning system for monitoring hazardous pollutants in coastal ecosystems and supports adaptive management strategies to mitigate biodiversity loss in urbanized seascapes.

## Introduction

Marine biodiversity is declining at an unprecedented rate due to the cumulative impacts of climate change, pollution, and coastal development (McCauley et al., 2015; O’Hara et al., 2021). Estuarine and nearshore ecosystems are particularly vulnerable, as they experience compounding pressures that continue to cause declines in ecosystem functioning and biodiversity (Lotze et al., 2006; Preston et al., 2025). In response, the Kunming–Montreal Global Biodiversity Framework (GBF) set ambitious targets to halt biodiversity loss, including expanding globally protected areas from 10% (Aichi Target 11) to 30% of the planet (Target 3). Despite global commitments to the GBF, the current network of marine protected areas (MPAs) only protects ∼7.5% of existing marine habitats (Arneth et al., 2023), with less than half of the existing MPAs fully established or managed (Aminian-Biquet et al., 2024; Gill et al., 2017). The low protection reflects the challenges of routine monitoring in marine environments and our comparatively limited understanding of the ecological dynamics shaping species distributions relative to other ecosystems (Herbert-Read et al., 2022; Ramírez et al., 2022). To further promote the establishment of new MPAs and facilitate adaptive conservation management strategies, effective and scalable tools for characterizing species distributions and community dynamics are urgently needed.

Environmental DNA (eDNA) metabarcoding presents a promising solution to conventional marine biodiversity assessment (Deiner et al., 2017; Djurhuus et al., 2020; Miya, 2022; Si et al., 2025). eDNA is DNA derived from environmental samples (i.e., seawater) that originates from tissue or cellular fragments discarded by the targeted organisms (e.g., skin shedding, faeces, or body fluids). Combined with high-throughput sequencing (e.g., eDNA metabarcoding), eDNA provides a rapid and non-invasive means to assess broad biodiversity dynamics, including elusive or cryptic marine species (Si et al., 2025). Additionally, eDNA-based biodiversity assessment is effective in dynamic, turbid, or remote marine environments, making eDNA particularly valuable for estuarine and coastal systems, where traditional survey methods often fail to capture the full extent of biodiversity (Sigsgaard et al., 2020). As such, eDNA-based methods are effective for monitoring fine-scale temporal and spatial changes in community composition (Sevellec et al., 2021; Seymour et al., 2021), facilitate early detection of invasive species (Larson et al., 2020; Martino et al., 2025), habitat degradation rates, and shifts in biodiversity patterns under a changing environment (Yao et al., 2022). Advancements in eDNA metabarcoding combined with continual growth of taxonomic reference databases have facilitated increasing adoption of eDNA protocols as part of conservation management strategies (Goldberg et al., 2016). The integration in eDNA-derived biodiversity data with existing ecological frameworks for adaptive conservation management is at the forefront of eDNA and conservation research interest (Cai et al., 2025; Pichler et al., 2025; Stephenson et al., 2024).

Biodiversity data requires robust analytical frameworks to infer ecological processes (Bush et al., 2017; Cavender-Bares et al., 2022; Jetz et al., 2019). In the complex and interconnected marine ecosystems, species respond idiosyncratically to various interacting stressors, including abiotic, anthropogenic, and biotic factors (Crain et al., 2008; Halpern et al., 2008; Maxwell et al., 2013). Joint species distribution models (JSDMs) are advanced ecological models that simultaneously estimate species’ responses to environmental gradients and biotic associations (Ovaskainen et al., 2017; Pollock et al., 2014; Warton et al., 2015), distinguishing patterns driven by environmental filtering from those arising through ecological interactions (Clark et al., 2014; Pollock et al., 2014; Tikhonov et al., 2020). In marine systems, JSDMs have been used to show that environmental filtering dominates broad-scale community structure, with residual associations providing additional insight into co-occurrence patterns (Mao et al., 2026; J. Zhao et al., 2025). Specifically, JSDMs have shown North Sea fish-community assembly is driven by spatial processes linked to temperature gradients (Montanyès et al., 2023). Fundamentally, JSDMs utilize Bayesian statistics making them highly adaptable for assessing large datasets, such as those derived from eDNA, offering a powerful and mechanistic means of understanding of biodiversity spatiotemporal dynamics (Pichler & Hartig, 2021). JSDMs have been applied to reveal emergent biodiversity patterns, delineate multi-species conservation priorities, and inform the design of protected areas that better capture ecological complexity and species co-distributions (Norberg et al., 2019; Pollock et al., 2014). As such, JSDMs are becoming a reliable modeling framework for assisting conservation planning, particularly in systems experiencing overlapping stressors, where species-specific responses are often obscured by interacting environmental variation (Bohnett et al., 2024; Stephenson et al., 2024).

The marine environment of Hong Kong is one of the most ecologically stressed ecoregions in Southeast Asia, largely due to rapid urbanization and high anthropogenic influence. Such ecological stress is a major conservation concern for the two resident cetacean species: the Indo-Pacific humpback dolphin (*Sousa chinensis*) and Indo-Pacific finless porpoise (*Neophocaena phocaenoides*) (Liu et al., 2023), as well as several non-resident cetaceans that transition periodically through Hong Kong (Liu et al., 2022). Both resident species occupy unique ecological niches within Hong Kong waters, with *S. chinensis* inhabiting turbid, shallow waters (<20 m), while *N. phocaenoides* prefers comparably deeper water and generally avoids shoreline areas (Fang et al., 2022). Consequently, *S. chinensis* is predominantly found along the western edge of Hong Kong, where the Pearl River estuary creates a highly turbid zone along the western edge of Lantau Island (Chan & Karczmarski, 2024). In contrast, *N. phocaenoides* has been spotted widely across Hong Kong, though population densities are much higher in the southern waters, where the water quality and visibility are much greater (Jefferson & Moore, 2020).

Both *S. chinensis* and *N. phocaenoides* are listed as “Vulnerable” by the IUCN (Jefferson et al., 2017; Wang & Reeves, 2017) and listed as target conservation species by the local Hong Kong government and as Grade I national key protected animals by the Chinese central government (https://www.wwf.org.hk, accessed 2025). In response, Hong Kong has established eight marine protected areas (MPAs) and one marine reserve, collective covering ∼5% of its marine environment, with six MPAs located around the Lantau Island (https://www.afcd.gov.hk, accesses 2025). However, existing MPAs and strategies have a limited effect on cetacean population recovery, particularly *S. chinensis*, which was estimated to be >200 individuals 30 years ago, has declined to ∼60 individuals (Jefferson, 2018). While more difficult to monitor, *N. phocaenoides* is also expected to have experienced a population decline of approximately 33% over the past 20 years (W. Lin et al., 2019). The lack of population recovery is likely due to many factors, including insufficient MPA size, misidentify effective areas for MPA placement, vessel traffic, and noise pollution (Lang et al., 2025), continuing anthropogenic pressure from sewage and nutrient runoff (Geeraert et al., 2021), and land reclamation in preferred cetacean habitat (Chan & Karczmarski, 2024). These factors, along with fishing activities, also negatively impact cetacean prey availability (e.g., fish), further complicating the underlying dynamics leading to declines in cetacean populations (Lin et al., 2021).

Here, we integrate eDNA metabarcoding with JSDMs to investigate how abiotic conditions, trophic relationships, and anthropogenic pressures shape cetacean–fish co-occurrence patterns across an estuarine seascape. The objectives of this study were to: (a) characterize spatial and seasonal variation in niche use across cetacean and fish communities; (b) evaluate the influence of environmental and anthropogenic predictors on cetacean occurrence; and (c) disentangle the relative roles of species sorting and predator–prey dynamics in shaping cetacean spatial distributions. Our findings offer new insight into species-specific vulnerability to environmental change and highlight the potential of eDNA–JSDM frameworks to inform adaptive conservation strategies in data-limited, high-impact coastal systems.

## Material and Methods

### Study site and Sample collection

Water samples were collected across five marine ecoregions surrounding the entirety of Lantau Island, Hong Kong (Figure 1b), including the core MPAs designated for the conservation of resident cetaceans (Table 1). These regions: Northern (NLI), Eastern (ELI), Southeastern (SELI), Southern (SLI), and Southwestern Lantau Island (WLI) were selected to capture a gradient in cetacean presence and absence, ranging from historically high frequency areas for *S. chinensis* and *N. phocaenoides* to regions with limited or no recorded sightings (Hung, 2008; Jefferson et al., 2002). The sites show pronounced environmental heterogeneity: northern areas are influenced by estuarine outflow from the Pearl River, characterized by high turbidity and nutrient enrichment, while eastern sites receive oceanic inflow from the South China Sea, resulting in clearer, more saline waters (Yu et al., 2021). Variability in vessel traffic, including spanning ferry corridors and fishing zones, were included as a gradient of anthropogenic pressure (Lang et al., 2025).

**Figure 1.**
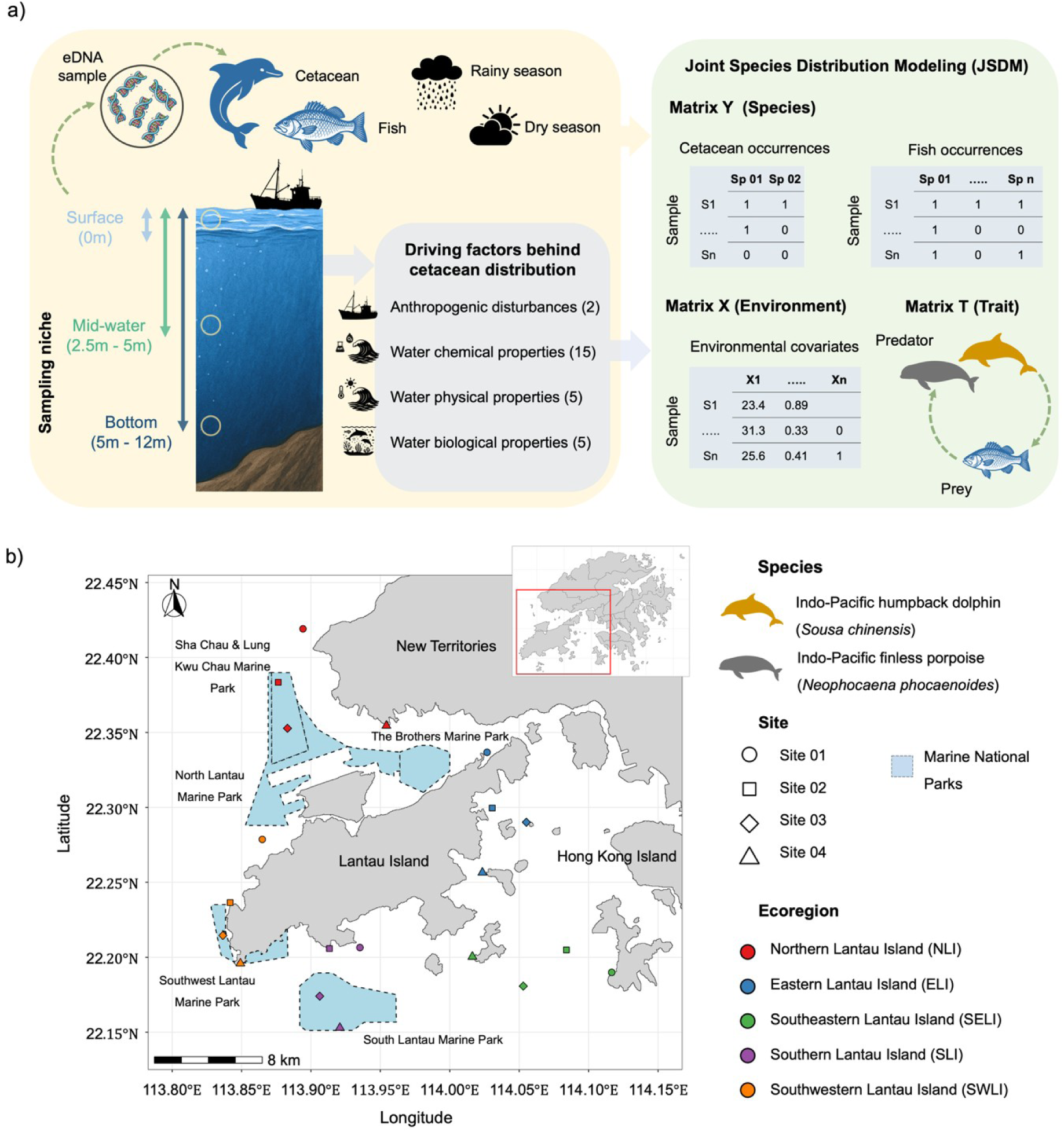
Overview of study design and sampling locations. (a) The analytical framework used to evaluate ecological and anthropogenic drivers influencing the distribution of resident cetaceans. (b) Map of sampling sites around Lantau Island, including locations situated within designated marine protected areas (MPAs) in Hong Kong.

**Table 1.** Summary of marine parks surrounding Lantau Island.

| Marine Park | Established date | Location | Area | Specific purpose |
| --- | --- | --- | --- | --- |
| Sha Chau and Lung Kwu Chau Marine Park | November 1996 | Open waters of western side of Hong Kong | 1,200 ha | - |
| The Brothers Marine Park | December 2016 | Northern Lantau waters | 970 ha | To help better conserve the Chinese White Dolphins their habitats and enhance the marine and fisheries resources therein. |
| Southwest Lantau Marine Park | April 2020 | Southwestern coastal waters adjacent to the Lantau. | 650 ha |  |
| South Lantau Marine Park | June 2022 | Southern Lantau waters | 2,067 ha | To help better conserve the Chinese White Dolphins and Finless Porpoise, their habitats and to enhance the fisheries resources therein. |
| North Lantau Marine Park | November 2024 | Northern Lantau waters | 2,400 ha | To better conservation of Chinese White Dolphins, their habitat, the marine ecology and fisheries resources in the area |

Overview of marine protected areas (MPAs) in the Lantau Island region, detailing their geographic location, surface area, and officially designated conservation objectives.

Environmental DNA sampling was conducted during the late rainy season (11-23 September 2022) and dry season (5-17 February 2023) to capture seasonal variability. Within each of the 5 ecoregions, 4 sites were sampled (20 sampling sites). At each site, triplicate 1 L seawater samples were collected using a depth sampler, collecting seawater at three depths: surface (0 m), mid-water (2-5 m), and near-bottom (5–12 m), adjusted according to local bathymetry (Figure 1a). To avoid cross-contamination, one depth sampler was used per sampling location. Prior to use, depth samplers were cleaned, dried, and decontaminated for DNA using DNA*Zap*^TM^ solution (Thermo Fisher Scientific Inc. USA). Sampling gear was prepared for each site individually before sampling and stored in separate sampling kit containers. All samples (N = 360) were filtered on-site aboard the sampling vessel using a field peristaltic pump (Geotech, Denver, USA) with sterile silicone tubing using enclosed 0.22 µm Sterivex filters (Millipore, USA). Field negative controls were processed for each site, consisting of autoclaved Milli-Q water (1 L per site) (n = 40). All filters were immediately sealed in clean bags, kept on ice during transport, and stored at –20°C upon arrival in the laboratory pending DNA extraction.

### Analysis of water quality (physical, chemical and biological properties) and anthropogenic disturbance measures

Twenty-four quality parameters (Supplementary Table S1) were assessed using nearby water monitoring stations with monthly data provided by the Environmental Protection Agency (EPA). Each eDNA sampling site was assigned to the nearest station within the same ecoregion (distance: 1–5 km). Environmental variables were extracted for the corresponding sampling period and matched depth strata (surface, mid-water, bottom) to ensure direct comparability with eDNA samples. Anthropogenic pressure was quantified using two AIS-derived (Automatic Identification System) layers from Global Fishing Watch (GFW; https://globalfishingwatch.org/): vessel presence and apparent fishing effort, representing general vessel activity and commercial fishing intensity, respectively. Data were extracted for the wet season (September 2022) and dry season (February 2023) (Supplementary Table S2). Vessel presence (hours per 0.5 km²) was calculated by the GFW as the cumulative hourly position of AIS-transmitting vessels within each grid cell. Apparent fishing effort (hours per 0.5 km²) was derived from GWS behavioural-classification algorithm, which identifies fishing activity based on characteristic movement patterns (e.g., speed and turning angle) of vessels classified as fishing vessels. Each AIS transmission is assigned a probability of fishing, and these values aggregated to generate spatial fishing-effort layers.

### Molecular analysis

All eDNA extractions and library preparations were performed in a designated and clean eDNA laboratory at the eDNA & eEcology lab at The University of Hong Kong. Operators strictly followed lab protocol to minimize contamination risks, including surface decontamination with 10% bleach and 70% ethanol, UV treatment, and regular cleaning of laboratory equipment and workspaces with DNA*Zap*^TM^ solution (ThermonFisher). Environmental DNA from seawater and field negative filters was extracted using QIAGEN DNeasy blood & tissue kit (Qiagen, Venlo, The Netherlands), following an established eDNA extraction protocol (Spens et al., 2017). The extracts were further purified using QIAGEN PowerClean Kit, quantified with a NanoDrop spectrophotometer (Thermo Fisher), and stored at –20 °C until library preparation.

A two-step PCR protocol was used to generate metabarcoding amplicon libraries (Bohmann et al., 2022; Seymour et al., 2020) for fish and cetacean communities separately, with the assistance of a Biomek i5 Automated Workstation (Beckman Coulter, USA). For the fish community, PCR1 (i.e., barcode amplification) was carried out using MiFish-U primers (Miya et al., 2020), while for cetacean communities, the ecoPrimers set was used (Riaz et al., 2011); both primer sets target different subsections of the mitochondrial 12S rRNA gene. PCR1 amplification was run in triplicate for each sample to increase overall library biodiversity yield. For the Mifish-U primers, the PCR1 reaction consisted of a total volume of 12 μl, including 6 μl of 2X KAPA HiFi HotStart ReadyMix (Roche, Switzerland), 0.4μl of each forward and reverse primer (10μM), 4.2 μl of ddH_2_O, and 1μl of template DNA. The PCR thermocycling program is as follows: an initial denaturation of 3 min at 95°C, followed by 35 cycles of 98°C for 20s, 65°C for 15s, and 72°C for 15s, with a final extension of 72°C for 5 min. For ecoPrimers, PCR1 reaction consisted of a total volume of 25 μl, including 12.5 μl of KAPA HiFi HotStart ReadyMix (Roche, Switzerland), 0.3 μl of each forward and reverse primer (10μM), 6.9 μl ddH_2_O, and 5 μl of template DNA. Thermocycler conditions are as follows: an initial denaturation at 95°C for 3 min, followed by 35 cycles of 98°C for 20s, 55°C for 30s, and 72°C for 15s, and a final extension of 72°C for 1 min. After pooling the PCR1 products triplicate, the pooled products were cleaned using AMPure XP beads (Beckman). PCR2 reactions consisted of a total volume of 25 μl with 12.5 μl KAPA HiFi HotStart ReadyMix (Roche, Switzerland), 5 μl Nextera UD indices (Illumina) and, 7.5 μl PCR1 template. Thermocycler conditions for PCR2 include an initial denaturation at 95°C for 3min, followed by 8 cycles of 98°C for 30s, 55°C for 30s, and 72°C for 30s, and a final extension of 72°C for 5 min. PCR2 products were then cleaned again using AMPure XP bead (Beckman). The PCR products, including the samples, field negatives, and PCR negative controls, were quantified using Qubit fluorescence (Thermo), normalized, and pooled in equimolar quantities in-house. The fish and cetacean libraries were then sent to the Centre for PanorOmics Sciences (CPOS) Genomics Core, The University of Hong Kong, and sequenced separately on Illumina Miseq platform (Illumina, USA) with 2 × 300 bp paired-end reads.

### Bioinformatics

Bioinformatics processing was carried out using a series of Linux-based scripts. First, primer sequences were removed using Cutadapt (Martin, 2011). Next, quality filtering was conducted using the *fastq_filter* command in VSEARCH (Rognes et al., 2016). Reads were then combined and dereplicated using the *derep_fulllength* function in VSEARCH, followed by denoising using the unoise3 algorithm implemented in VSEARCH. Lastly, chimera removal was performed using the *uchime3_denovo* function in VSEARCH. The resulting Amplicon Sequence Variant (ASV) frequency table was then used for subsequent taxonomy assignments and statistical analyses.

BLAST+ was used to assign taxonomic information to each of the unique Amplicon sequence variants (ASVs) created in the prior bioinformatics step, checked against a locally curated database of NCBI sequence data. The local database was generated using the CRABS workflow, with a separate databased generated for each of the primer sets (Jeunen et al., 2023). The BLAST+ command was carried out using a Linux-based script. The output file was then closely examined for each ASV taxonomic identification. All ASVs were excluded that did not have taxonomic identification to at least the domain level or were not identified as marine species. The dataset was screened with a threshold of 97% percentage identification for species, 90% for genera, and 80% for family. ASVs were filtered for contamination based on their read counts in the negative controls, whereby the maximum value of any ASV detected across the negative controls was removed from all samples ASV pools.

### Statistical analysis

All analyses were conducted in R version 4.2.1 (R Core Team, 2022). Prior to downstream analysis, we assessed the effects of spatiotemporal patterns and sample depth on cetacean richnesss and relative abundance using the full dataset. We found that depth was not associated with changes in cetacean richness or relative abundance across the full cetacean amplicon dataset (Appendix S1). Therefore we restricted subsequent analyses comparing cetacean and fish biodiversity dynamics to over lapping samples, which including all middle depth samples from all sites and seasons. This restriction was not a removal of data given there was no loss in the observed richess or relative abundance. We provide the initial check of our eDNA sampling methods prior to matching the sampling effort with the target trophic fish group (Supplementary Appendix 1). To standardize sampling effort, triplicate samples for each combination of season, ecoregion, site, and depth were pooled into a single composite sample by taking the mean proportional reads across each replicate set of within site samples.

To assess sampling sufficiency across depths and seasons, we generated sample-size-based rarefaction and extrapolation (R/E) curves for sample completeness and species richness for both cetaceans and marine fish collected from eDNA samples. The sample completeness R/E curves assess the adequacy of the sampling through sample coverage, which represents the proportion of the Species richness (Hill number *q* = 0), was estimated using the “iNEXT” package (Hsieh et al., 2016).

To test the effects of season and depth on species richness and alpha diversity (Shannon diversity, H′, and Pielou’s J evenness) of cetaceans and fish, we fitted generalized linear mixed models (GLMMs). Species richness was modelled using a negative binomial error distribution.

Shannon diversity was modelled using a Tweedie distribution with a log link, while evenness was modelled using a beta distribution with a zero-inflation term to accommodate bounded continuous values with excess zeros. Season and depth were included as fixed effects, and sampling ecoregion around the island was included as a random intercept. Species richness models were fitted using the *glmer.nb* function in the *“*lme4” package, whereas models for Shannon diversity and evenness were fitted using the *glmmTMB* function in the “glmmTMB” package.(Magnusson et al., 2020).

To assess patterns in community composition across seasons and depths, non-metric multidimensional scaling (NMDS) based on Bray–Curtis dissimilarities using the *metaMDS* function from the “vegan” in R (Oksanen et al., 2022) was used. Whether the season and the depth change the cetacean and fish community composition was assessed using permutational multivariate analysis of variance (PERMANOVA) using Bray–Curtis dissimilarity with 999 permutations using the *adonis2* function available in the “vegan” package (Oksanen et al., 2022). A pairwise comparison was conducted to test the significance of each season’s depth *pairwise.adonis* function available in the “pairwiseAdonis” package (Martinez Arbizu, 2020). The Mantel test was conducted to evaluate the correlation between fish and cetacean abundance-based dissimilarity matrices using Spearman’s rank correlation, using 9,999 permutations to assess statistical significance.

To assess the effects of ecoregion and season on the relative abundance of resident cetaceans, we fitted GLMMs using the “glmmTMB” package with a Tweedie distribution with a log link, including sampling site as a random intercept. The Tweedie index parameter (*p*) was estimated by maximum likelihood and verified to fall within the compound Poisson–Gamma range (1 < *p* < 2) (See Appendix S2 for details). Estimated marginal means and pairwise comparisons were computed using the “emmeans” package to evaluate spatiotemporal differences (Lenth, 2023).

The collected water quality and anthropogenic disturbance factors were used to assess their impact on the relative abundance of resident cetaceans. Prior to modeling, multicollinearity among predictors was evaluated using the “linKET” package (Huang, 2021). A Spearman correlation matrix was first calculated, and highly correlated predictyors (|r| ≥ 0.7) were identified. Variables with a variance inflation factor (VIF) ≥ 5 were considered highly correlated and were then removed to reduce the collinearity (van Steenderen & Sutton, 2024). All remaining predictors showed acceptable level of multicollinearity prior to model fitting.

We fitted a GLMM using a Tweedie distribution with a log link to examine the environmental drivers of resident cetacean relative abundance. For each cetacean species, relative abundance at each site was calculated by dividing site-specific read counts by the species-specific total number of reads, providing a semi-quantitative proxy of relative signal intensity rather than absolute abundance. Although read abundance does not represent absolute population size, previous studies have shown that eDNA read counts are positively associated with organismal abundance or biomass in aquatic systems, providing a defensible semi-quantitative proxy for relative abundance at local scales (Lacoursière-Roussel et al., 2016; Nakagawa et al., 2022). Here, the relative abundance of *S. chinensis* and *N. phocaenoides* was used as the response variable, and the filtered environmental predictors and fixed factors with the sampling ecoregions included as a random factor to control for potential confounding. No residual over dispersion was detected in any final GLMM models. The model fitting was performed using the *glmmTMB* function from the “glmTMB” package.

To disentangle the effect of species sorting (determined by the effects of environmental filtering) and predator–prey interactions (cetacean-fish association), we used Joint Species Distribution Modeling (JSDM) with the Hierarchical Modeling of Species Communities (HMSC) framework using the “HMSC” package available in R (Tikhonov et al., 2020). HMSC simultaneously estimates responses of species to environmental conditions and residual associations attributable to biotic interactions while accommodating hierarchical sampling structures (Ovaskainen et al., 2017). Within the framework, cetaceans and fish were modelled jointly as response variables, with cetaceans treated as predator species and fish as their prey. Species occurrences and relative abundance derived from eDNA metabarcoding were used as response variables, with environmental predictors treated as fixed effects and spatial random effects included at the ecoregion level to capture unmeasured variation. All environmental predictors were standardized prior to modeling using the z-score transformation.

We fitted two hurdle models, each comprising a binomial model for presence–absence and a log-normal model for relative abundance, to account for zero inflation in the eDNA data (56.85% and 89.87% for cetaceans and fish, respectively). We used the log-normal component to model continuous variation in relative abundance. Although not explicitly designed for proportional data, this approach is a practical choice when eDNA read data are interpreted as semi-quantitative indicators of relative abundance, as supported by previous evaluations of fish eDNA abundance signals (Odriozola et al., 2021; Rourke et al., 2022). The null model included only the intercept, while the full model incorporated filtered environmental predictors. Random effects were structured at each ecoregion level. Residual associations between species were estimated using a latent factor model within HMSC. We specified a prior range of five to ten latent factors (nfMin = 5, nfMax = 10) for the spatial level, allowing the model to adaptively determine the effective number of factors during MCMC sampling (adaptNf). The latent factors induce a species-to-species residual covariance matrix (ω), which represents unexplained co-variation after accounting for environmental predictors and hierarchical structure. Species association matrices were extracted from the posterior mean correlation matrices for both the null and full models. Significant positive and negative associations were identified based on a posterior support threshold of 0.75. Association in the null model reflect shared environmental preferences and unmodelled structure, whereas those in the full model matrix represent residual associations after accounting for environmental predictors and were interpreted as putative biotic interactions. Variance partitioning was conducted to quantify the contributions of environmental predictors and spatial random effects to total explained variance. Association matrices were visualized using “corrplot” package (Wei et al., 2017) and variance components were plotted using “ggplot2”.

Model performance was assessed using the area under the curve (AUC) for presence–absence and *R²* for relative abundance data. We further estimated (1) explanatory power, using the same dataset used to fit the model, and (2) predictive power through four-fold cross-validation. Within the predictive framework, we distinguished between (a) unconditional predictions, standard cross-validated predictions without species conditioning, and (b) conditional predictions, which incorporated information on species associations from the training set to improve forecasts for co-distributed taxa. By jointly modeling the community data, HMSC quantifies the relative contributions of environmental filtering and biotic associations to observed cetacean community patterns.

The HMSC models were fitted using Markov Chain Monte Carlo (MCMC) with default priors. We ran four independent chains, each consisting of 37,500 iterations. The first 12,500 iterations were discarded as a burn-in period, and the remaining samples were thinned by 100, resulting in 250 posterior samples per chain, totaling 1,000 samples across all chains. Finally, we evaluated the convergence of the MCMC model by calculating the effective sample size and the potential scale reduction factor (See Supplementary S2 for details). We evaluated sensitivity to latent factor dimensionality by refitting the matched-subset HMSC model with the number of latent factors fixed across a grid (nf = 1-4), while keeping the fixed-effects structure identical across models. Predictive performance was stable across values of nf, and inferred residual cetacean–fish associations remained weak, indicating that results are not driven by latent over-parameterization.

## Results

12S ecoPrimer amplicon libraries included an initial 400 libraries (360 eDNA samples and 40 blanks), with 368 successfully sequenced, resulting in 117,602,658 12S (average 307,860 per sample) single reads. Successful libraries included 165 rainy season eDNA samples and 167 dry season eDNA samples. With 118 libraries from mid-depth, 110 from surface water and 104 from bottom depth (Supplementary Figure S1). Due to logistics constraints and confirmed lack of depth influence on the wider dataset (Supplmentary Appendix 1), 12S Mifish-U amplicon libraries including an initial 180 libraries (155 eDNA samples and 25 blanks), with 179 successfully sequenced. Successful libraries included 120 mid-depth, 19 surface water and 15 bottom depth samples (Supplementary Figure S2). The Mifish-U libraries generated 51,395,206 (average 273,379 per sample) single reads.

Please note that cetacean–fish co-occurrence analyses was conducted on samples that overlapped in their sequencing effort to ensure that cetacean and fish detection, and potential interaction, originated from the same sampling events. eDNA replicate samples collected within each sampling event location and depth were reduced to a single composite sample to reduce stochastic detection and sequencing noise, which can be driven by PCR and sequencing rather than ecologically based (Deiner *et al*. 2017, Seymour *et al*. 2021). As such, detection probability is not explicitly modeled here, however the conservative reduction improves the overall robustness of community-level estimates. Because depth coverage in the MiFish-U subset was uneven, co-occurrence analyses were conducted on the full matched dataset rather than stratified by depth to preserve sample size and statistical power. While this sample size is modest, previous studies have shown that JSDM performance depends not only on sample size but also on model structure and species-specific characteristics, and increasing sample size does not necessarily improve predictive performance across all models or species (Xu et al., 2025)

Sampling coverage indicated that mid-water samples consistently achieved the highest completeness for both cetaceans and fish, although coverage for fish assemblages remained lower overall (Supplementary Figure S3). Species richness (q = 0) for cetaceans approached asymptotic levels across all depths during the rainy season, and nearly reached asymptote in mid-water samples during the dry season. In contrast, fish species richness did not reach an asymptote for some depths. As such fish community detection may be incomplete, and therefore inferences regarding cetacean–fish associations should be interpreted with appropriate caution.

### Seasonal shifts in fish communities but not cetaceans

Cetacean species richness (8 species in total) was significantly affected by season (χ² = 11.92, *df* = 1, *p* < 0.001), whereas Shannon diversity showed a marginal effect of depth, and evenness was not significantly influenced by either factor. In contrast, a total of 199 fish species found and varied between seasons (rainy season: 15.97 ± 7.47; dry season: 21.56 ± 8.75) and among locations (NLI: 23.54 ± 9.03; ELI: 19.44 ± 8.52; SELI: 16.31 ± 8.74; SLI: 20.05 ± 8.41; WLI: 15.55 ± 6.76). Fish species richness was marginally influenced by the interaction between season and depth (χ² = 4.95, *df* = 2, *p* = 0.084). Fish Shannon diversity was strongly affected by season (χ² = 15.67, *df* = 1, *p* < 0.001), while fish evenness varied significantly with both season (χ² = 23.26, *df* = 1, *p* < 0.001) and depth (χ² = 7.97, df = 2, *p* = 0.019) (Figure 2).

**Figure 2.**
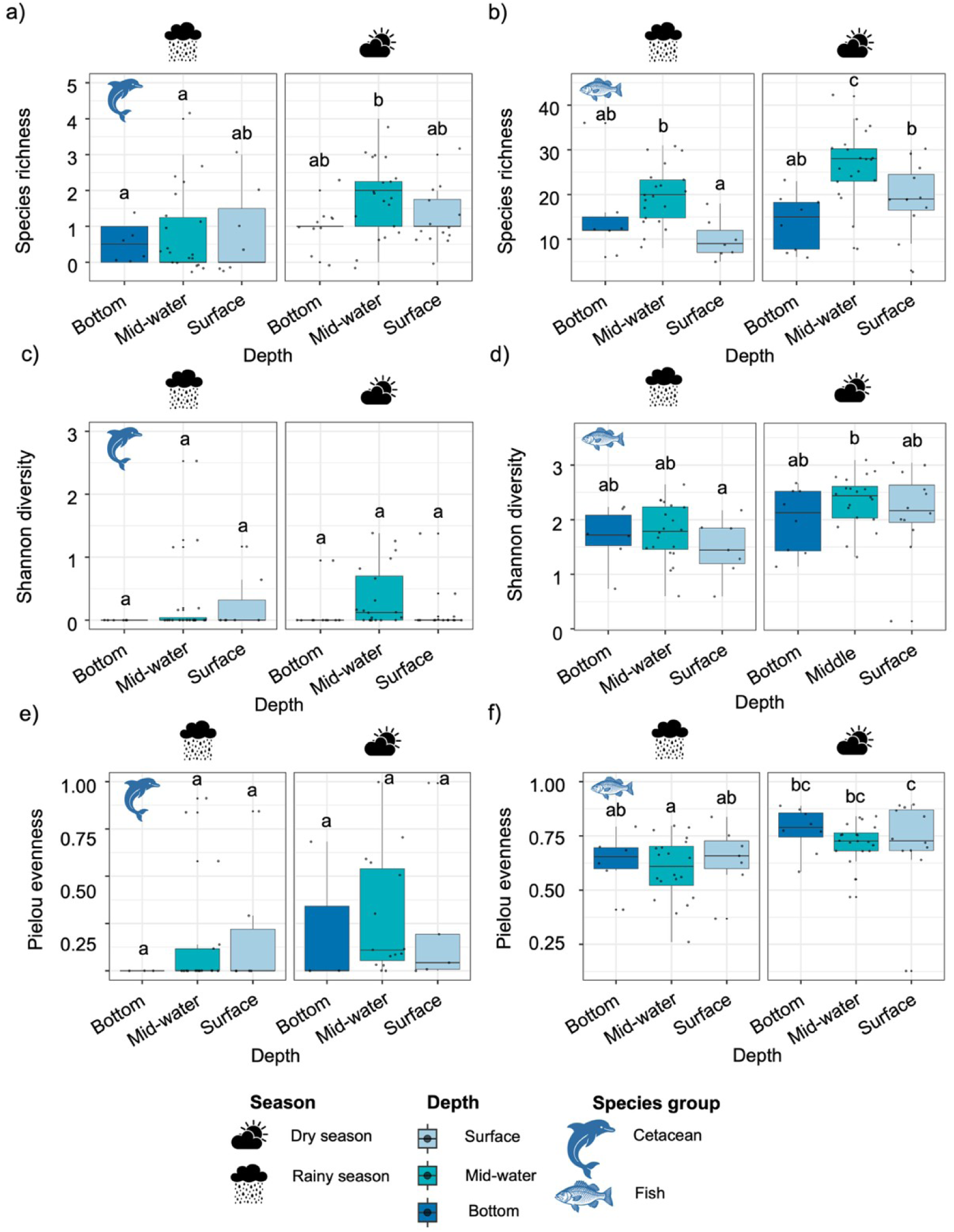
Seasonal variation in alpha diversity of fish and cetacean communities. The alpha diversity metrics of the fish (left column) and dolphin (right column) across rainy and dry seasons. Species richness shows the total number of species detected; Shannon diversity (H′) reflects both richness and evenness; and Pielou’s evenness (J) quantifies community evenness. The compact letters denote statistically significant differences in diversity metrics across seasons and depths (*p* < 0.05).

NMDS ordination revealed no clear clustering of cetacean communities by season or depth, with weak community separation (stress = 0.1378; non-metric fit *R*^2^ = 0.981; linear fit *R*^2^ = 0.911) (Figure 3a). In contrast, fish community composition showed pronounced seasonal separation (stress = 0.1763; non-metric fit *R*^2^ = 0.969; linear fit *R*^2^ = 0.85) (Figure 3b; see also occurrence-based NMDS in Supplementary Figure S4), although depth-related clustering was less distinct.

**Figure 3.**
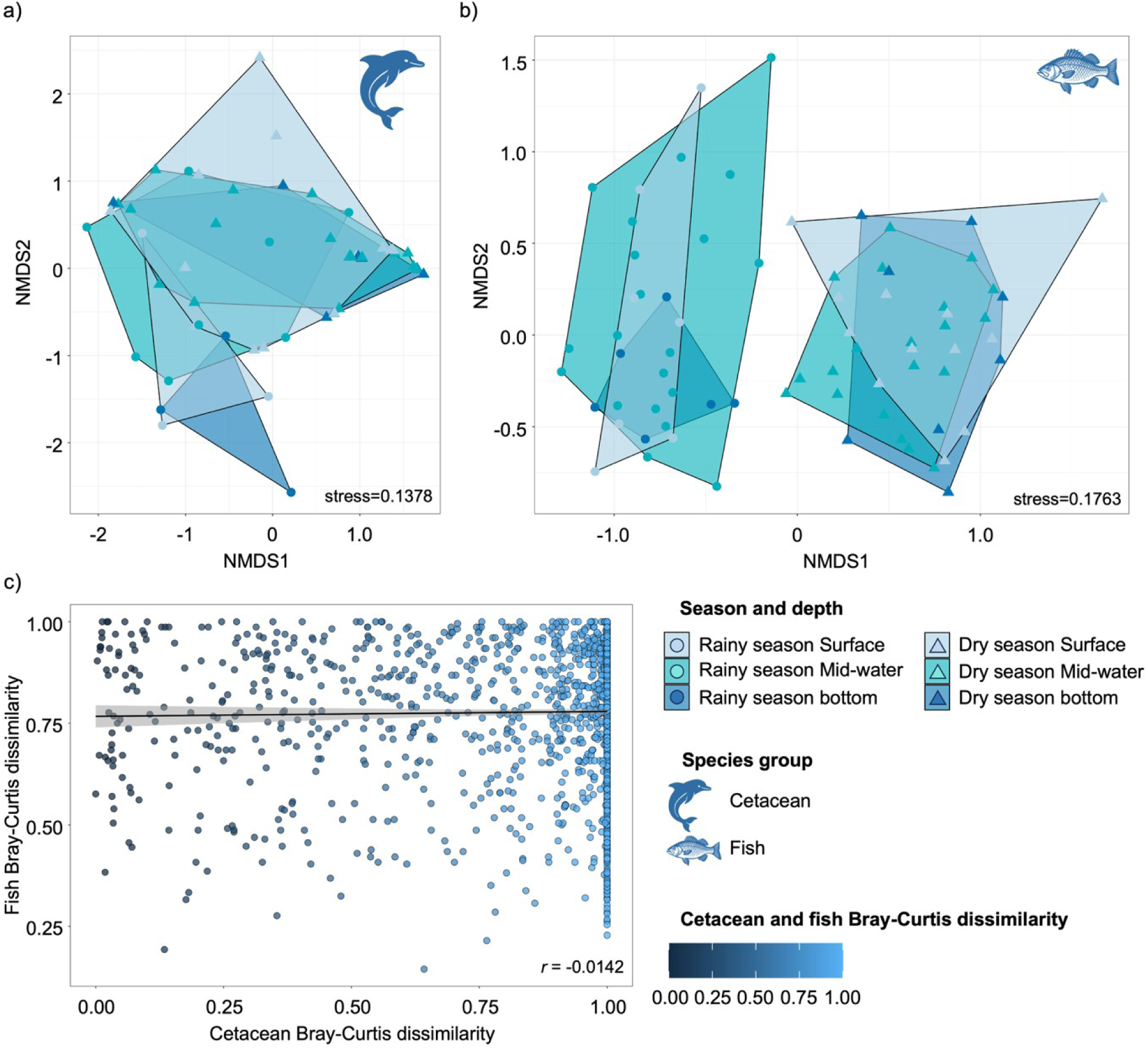
Seasonal and vertical structuring of marine cetaceans and fish. The community composition of (a) cetaceans and (b) fish community changes over season (rainy vs dry) in different depth strata (surface, mid, bottom) in marine environment using nonmetric multidimensional scaling (NMDS) based on Bray–Curtis dissimilarity. Cetacean and fish Bray-Curtis dissimilarity showed (c) weak negative correlation between fish and cetacean community changes.

PERMANOVA results confirmed these patterns: fish community composition varied significantly by season–depth interaction (pseudo-*F_1,5_*= 3.94, *R*² = 0.23, *p* = 0.001), while no significant patterns were observed in cetaceans (Supplementary Table S3). Post hoc pairwise comparisons showed that fish community composition between seasons was significantly different, but not between depths at each season (Supplementary Table S4). PERMDISP indicated that differences in fish composition were driven by deterministic turnover rather than differences in dispersion (*F_5,72_* = 1.646; *p* = 0.156). The Mantel test showed no significant correlation between cetacean and fish community dissimilarity matrices (*r* = −0.0142, *p* = 0.631).

### Spatial variation in resident cetacean abundance and environmental predictors

Generalized linear mixed models showed that the relative abundance of *S. chinensis* varied significantly by season (χ² = 7.79, df = 1, *p* = 0.005) and ecoregion (χ² = 62.30, df = 4, *p* < 0.001), with the highest relative abundance recorded in SWLI during the dry season (Figure 4a). No detections were recorded in SELI across seasons. *N. phocaenoides* abundance also varied significantly by season (χ² = 37.24, df = 1, *p* < 0.001) and ecoregion (χ² = 51.75, df = 4, *p* < 0.001), with peak abundance in SELI, SLI, and SWLI during the dry season (Figure 4b and Supplementary Figure S5).

**Figure 4.**
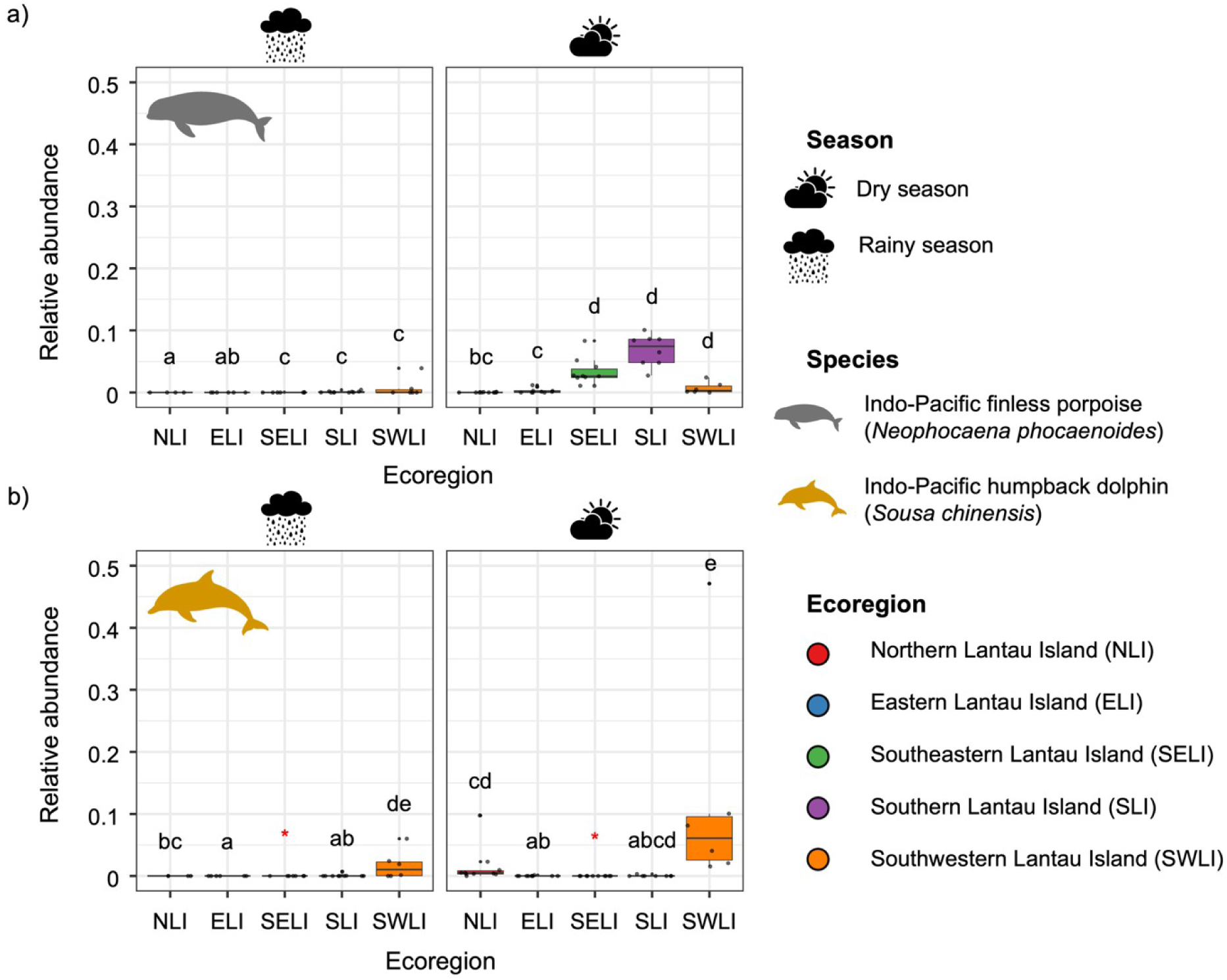
Spatiotemporal variation in relative abundance of resident cetaceans. Relative abundance of (a) Indo-Pacific finless porpoises (*Neophocaena phocaenoides*) and (b) Indo-Pacific humpback dolphins (*Sousa chinensis*) during the rainy and dry seasons. Colored asterisks (*) mark ecoregions where humpback dolphins were not detected in either season or were excluded from statistical comparisons. Compact letters indicate significant spatiotemporal differences in relative abundance (*p* < 0.05).

After removing the collinear predictors, seven environmental predictors were retained for modeling (Supplementary Figure S6 and S7). For *S. chinensis,* Secchi disc depth (m) (*β* = -7.23, *p* < 0.001), and presence of vessels (hours/0.5 km²) (*β* = -0.006, *p* = 0.002) were significantly negatively associated with relative abundance, whereas nitrate nitrogen (mg/L) showed significant positive effect (*β* = 14.26, *p* < 0.001). In contrast, *N. phocaenoides* showed a significant negative association with the nitrate nitrogen (mg/L) (*β* = -13.47, *p* < 0.001) effect and marginally significant negative effect with E. coli concentration (CFU/100 mL) (*β* = -0.21, *p* = 0.086).

### Species sorting dominates cetacean–fish co-occurrence patterns

Diagnostic metrics indicated good MCMC convergence for both HMSC models (Supplementary Figure S8). Mean effective sample sizes were high for both models, with *β* parameters (species sorting) averaging (mean ± SD= 1052.28 ± 133.64) in the null model and (mean ± SD=1047.98 ± 133.50) in the full model, and ω (latent factor) parameters averaging (mean ± SD=1039.69 ± 136.95) and (mean ± SD=1041.17 ± 125.83) respectively. Potential scale reduction factors (PSRF) were consistently close to 1, with mean ± SD of (mean ± SD=1.01 ± 0.06) (*β*) and (mean ± SD=1.05 ± 0.06) (ω) in the null model, and (mean ± SD=1.01 ± 0.01) (*β*) and (mean ± SD=1.03 ± 0.03) (ω) in the full model (Supplementary Figure S9), confirming adequate mixing and convergence of the MCMC chains. Stable predictive performance (AUC/R²) and explanatory power across nf = 1-4 confirm that species association inferences are robust and not sensitive to latent factor dimensionality (Supplementary Figure S10).

In both the binomial and log-normal models, the species association matrices for the null model showed weak positive and negative co-occurrence patterns between cetaceans and individual fish species, including known prey taxa (Supplementary Figure S11a). However, these associations disappeared in the full model after accounting for environmental covariates (Supplementary Figure S11b), suggesting that shared environmental responses explained more of the observed co-occurrence structure than detectable residual predator–prey associations at the spatial and temporal resolution.

Explanatory power for species occurrences was consistently higher in the full models compared to the null models, for both fish and resident cetaceans (Table 2). In the binomial models, predictive performance was generally stronger for cetaceans than for fish, particularly when environmental covariates were included. For fish, the inclusion of environmental variables improved occurrence-based predictive power (AUC) by 0.216 units (from 0.246 to 0.462), whereas accounting for cetacean presence did not enhance predictive power for fish. In contrast, the predictive power for cetacean occurrences increased markedly with environmental covariates (from 0.419 to 0.668). The addition of co-occurrence information further improved performance, suggesting that environmental filtering played a dominant role in shaping species distributions. For log-normal models, predictive power remained low overall. Fish abundance models showed negative cross-validated R² values (−0.425 to −0.085), indicating poor transferability to held-out data. Accordingly, abundance-based inferences, including residual species associations, should be interpreted with caution. In contrast, cetaceans occurrence-based models showed improved predictive performance (R² = −0.083 to 0.039) with environmental covariates, providing stronger support for environmentally structured distributions than for abundance-based species associations.

**Table 2.** Summary of joint species distribution model performance for resident cetaceans and fish communities.

| Category/<br>Measure | Species<br>group | Model | No. of<br>species | Explanatory<br>power | Unconditional<br>predictive<br>power | Conditional<br>predictive<br>power |
| --- | --- | --- | --- | --- | --- | --- |
| Occurrences<br>/ AUC | Fish | Null | 199 | 0.639 | 0.246 | 0.241 |
|  |  | Full | 199 | 0.815 | 0.462 | 0.458 |
|  | Cetacean | Null | 2 | 0.685 | 0.419 | 0.426 |
|  |  | Full | 2 | 0.784 | 0.668 | 0.696 |
| Abundance/<br>R<sup>2</sup> | Fish | Null | 199 | 0.296 | -0.425 | -0.480 |
|  |  | Full | 199 | 0.547 | -0.085 | -0.083 |
|  | Cetacean | Null | 2 | 0.225 | -0.083 | -0.083 |
|  |  | Full | 2 | 0.363 | 0.039 | -0.004 |

Explanatory, conditional, and unconditional predictive power of the HMSC framework applied to *S. chinensis*, *N. phocaenoides*, and associated fish species in the inshore waters of Lantau Island, Hong Kong SAR. *Explanatory power* quantifies how well the model fits observed data using all available covariates and random effects. *Unconditional predictive power* reflects the ability of the model to predict new observations under standard cross-validation. *Conditional predictive power* reflects prediction accuracy under species-wise conditional cross-validation, in which predictions incorporate cross-species association information through the residual covariance structure of the model.

Variance partitioning revealed distinct species-specific sensitivities to environmental and anthropogenic drivers (Figure 5 and Table 3). Anthropogenic disturbances explained the majority of the variance for *S. chinensis* occurrence (30.04%), while *N. phocaenoides* was primarily influenced by physical water properties, which accounted for 40.9% of explained variance (Figure 5a). In terms of relative abundance, both cetacean species were most affected by physical water properties (*S. chinensis*, 56.93%; *N. phocaenoides*, 52.77%). For the fish community, variance was more evenly distributed across predictor groups, with comparable contributions from anthropogenic disturbances, water chemistry, and physical conditions (Figure 5b). Spatial random effects at the ecoregion level explained only a small fraction of variance (≤9% for all taxa). This indicates that most variation was captured by measured environmental predictors. These differences reflect divergent habitat use and ecological niches between the two resident cetacean species.

**Figure 5.**
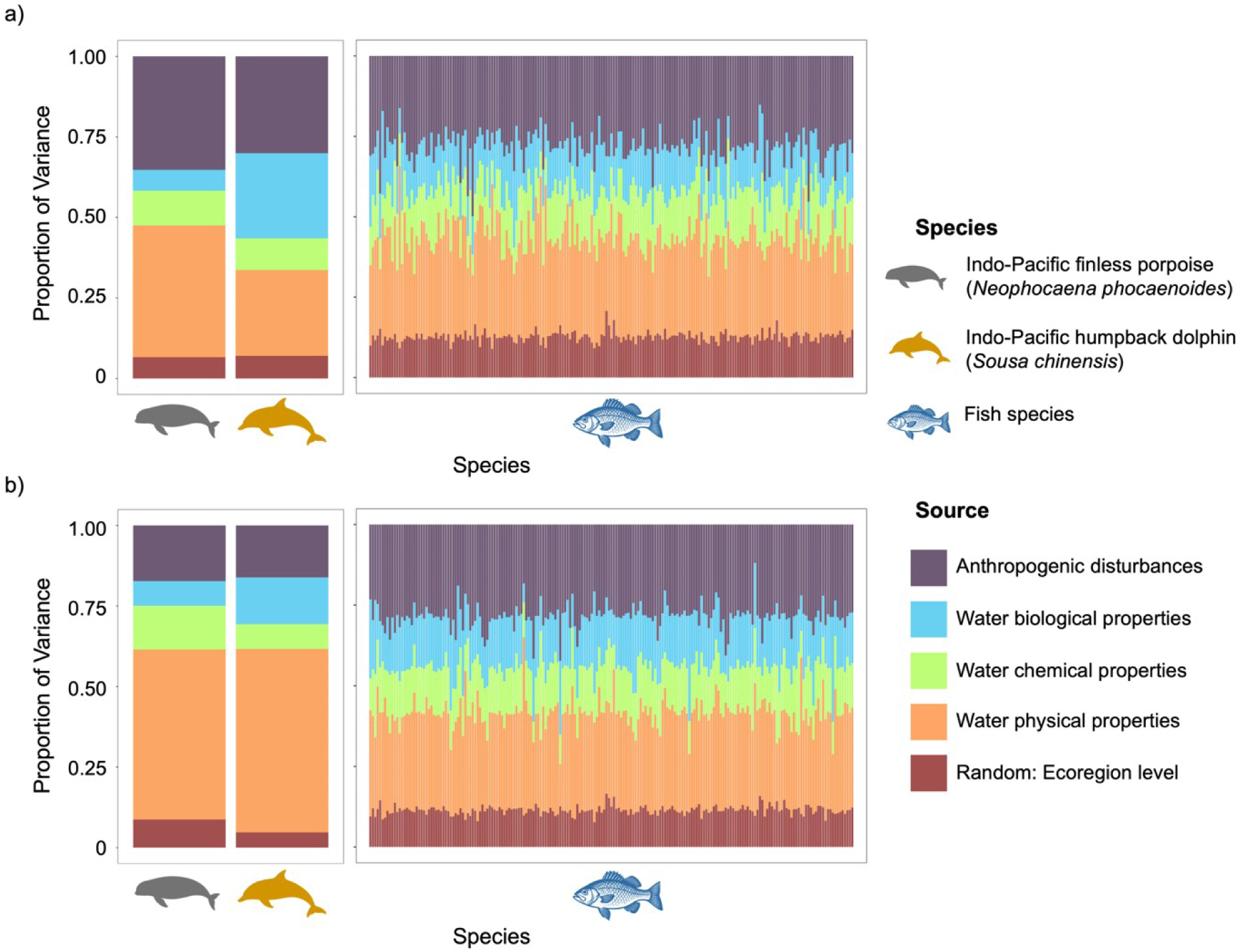
Variance partitioning of environmental drivers of community structure. Proportion of variance explained by environmental covariates and ecoregion-level random effects for (a) occurrence and (b) relative abundance data. Bars show the proportion of variance explained by grouped environmental variables from both binomial models.

**Table 3.** Partitioning of variance in species distributions for resident cetaceans and fish.

| Sources | Percentage of Variance (%) |  |  |  |  |  |
| --- | --- | --- | --- | --- | --- | --- |
|  | Predator |  |  |  | Prey |  |
|  | Indo-Pacific finless porpoise |  | Indo-Pacific humpback dolphin |  | Fish community |  |
|  | Occurrence based | Abundance based | Occurrence based | Abundance based | Occurrence based | Abundance based |
| Anthropogenic disturbances | 35.2 | 17.25 | 30.04 | 16.1 | 28.34 | 28.5 |
| Water biological properties | 6.48 | 7.66 | 26.44 | 14.52 | 13.76 | 15.69 |
| Water chemical properties | 10.77 | 13.57 | 9.86 | 7.69 | 14.16 | 14.19 |
| Water physical properties | 40.9 | 52.77 | 26.63 | 56.93 | 31.07 | 30.20 |
| Random: Ecoregion level | 6.63 | 8.74 | 7.02 | 4.76 | 12.66 | 11.4 |

Summary of mean variance contributions for each resident cetacean species and the overall fish community, as estimated by the JSMD model. Variance components reflect the relative influence of fixed and ecoregion level random effects.

## Discussion

This study presents the first spatially explicit assessment of cetacean distributions using an integrated framework that combines eDNA metabarcoding with JSDMs. By simultaneously modeling environmental conditions and prey availability, we disentangled the relative contributions of abiotic stressors and biotic associations in shaping the spatial patterns of two resident cetaceans in Hong Kong waters. Environmental DNA metabarcoding resolved fine-scale distributional patterns of individual cetacean species and their associated prey within one of the world’s busiest and heavily disturbed coastal systems. Applying JSDMs to eDNA data represents a significant advancement in applied eDNA research, providing robust insights and actionable recommendation for conservation planning and biodiversity management (Ovaskainen et al., 2017; Pollock et al., 2014).

### Contrasting spatiotemporal dynamics of cetaceans and fish

Cetacean species detection increased with mid-depth sampling and during the rainy season, with seasonal variation being relatively modest. This consistency reflects the dominance of two resident species and their relatively static distribution in Hong Kong waters. The limited temporal turnover between seasons indicates strong site fidelity and demographically constrained populations, reinforcing the urgency of local-scale threat mitigation to safeguard long-term persistence (Davidson et al., 2009). In contrast, fish communities showed seasonal shifts in species richness, diversity, and community composition. However, that fish richness did not reach an asymptope for most of the sampling sites indicates limit statistical power to detect weak species associations, and therefore non-detection should not be interpreted as evidence of absence. The observed seasonal variation in fish community composition may, cautiously, reflects recruitment pulses, ontogenetic habitat shifts, and other life-history processes (Nagelkerken et al., 2000). Despite these dynamics, evenness remained relatively stable, suggesting no single taxon dominated seasonally, potentially as a result of top-down dynamics (Saleem et al., 2012), though additional assessments are needed for full consideration.

### Environmental stressors shape resident cetacean distribution

Disentangling the effects of environment versus biotic interactions on species’ distribution is important for understanding local ecological dynamics need for developing effective conservation management policies (Wisz et al., 2013). Here we found that within the western waters of Hong Kong, cetacean distributions were more strongly associated with measured environmental covariates than with detectable residual biotic associations under the current sampling and modeling framework (i.e. fish communities). Hong Kong is densely populated, both locally and regionally as part of the greater bay area, resulting in a heavily disturbed coastal ecosystem. As such, habitat degradation, pollution and intense vessel traffic often overshadow trophic interactions within the region (Piwetz et al., 2021; Yuen et al., 2025). In contrast, under the current fish eDNA coverage and analytical framework, we did not detect a robust association between spatial variation in fish biodiversity and cetacean distribution patterns.

Prior studies using similar eDNA methods have indicated strong fish biodiversity responses to environment (Seymour et al., 2025) and trophic interactions (Boyse et al., 2025), highlighting the power of eDNA to capture fine scale community dynamics. Here we detected nearly 200 species of fish across the study area, capturing several key prey species of both cetacean species, including *Collichthys lucidus*, *Johnius belangerii* and *Clupanodon thrissa* (W. Lin et al., 2021; Zhang et al., 2023), but did not detect consistent temporal or spatial correspondence between fish community patterns and cetacean distributions under the current sampling and analytical framework. Additionally, the low predictive performance of the fish relative abundance component further limits confidence in relative abundance-based inference and reinforces our focus on occurrence-based patterns. Importantly, *Sousa chinensis* is known for being a highly opportunistic feeder, taking advantage of the most abundantly available prey when their preferred prey (e.g., *Larimichthys* spp. or *<u>Sardinella</u>* spp.) are unavailable (Lin et al., 2021). Prior assessments also discuss that neither bottom-up nor top-down control appear to dominate cetaceans-fish dynamics in Hong Kong waters, as degraded water quality and estuarine discharge, and heavy vessel traffic and coastal construction continuously shift the ecological landscape, potentially obscuring stable trophic relationships (Yuen et al., 2025).

After accounting for environmental covariates, no strong residual co-occurrence was detected, which may reflect limited statistical power and incomplete characterization of fish community structure in addition to ecological processes, suggesting that environmental predictors explain a greater proportion of the observed spatial variation than detectable residual associations at the resolution considered, with clear species-specific sensitivities. Relative abundance of *Sousa chinensis* declined with increasing Secchi depth (i.e., clearer water), consistent with well-documented preference for turbid, estuarine habitats characterized by shallow, sediment-rich coastal environments (Lin et al., 2021). In contrast, vessel presence showed a negative association with relative abundance, consistent with the disruptive effects of coastal anthropogenic activity, including noise and traffic (Todd et al., 2015). Although, *S. chinensis* are known to exhibit trawler-following behavior (Parsons, 2004), small fishing vessels, as common in Hong Kong are also elevates strikes risk (Kot et al., 2022; Slooten et al., 2013). Collectively, the patterns suggest that clearer waters and reduced vessel activities are key for population stability and recovery (Jefferson et al., 2023; Piwetz et al., 2021), consistent with increased sightings during COVID-19 when boating activity declined (Davidson H., 2020; Huang et al., 2024; Knott K., 2022). Spatially, *S. chinensis* was largely absent in southeast and eastern Lantau, likely due to chronic acoustic and physical disturbance, which disrupts foraging and social behaviour (Erbe et al., 2019; Pirotta et al., 2015). In contrast, higher relative abundance in western Lantau, particularly during the dry season suggests its role as a seasonal refuge, possibly linked to calving, prey shifts, or reduced human activity (Hung & Jefferson, 2004; Jefferson et al., 2012). Elevated sediment loads and vessel pressure near the Pearl River outlet further support concerns over cumulative stressors, which compromise habitat quality for *S. chinensis* (Erbe et al., 2019).

*Neophocaena phocaenoides* showed higher occurrence in the western and southern waters but were rarely detected in the most northern sites, near the Pearl River outlet, where elevated concentrations of nitrate nitrogen and E. coli suggest poor water quality. The presence of *N. phocaenoides* in heavily trafficked areas suggested that boating activity, while a significant effect, was not the main influence on their distribution. Instead, degraded water quality was the dominant factor shaping *N. phocaenoides* habitat use (Z. Xu et al., 2021). This divergence highlights how two ecologically similar coastal cetaceans can exhibit distinct vulnerabilities: *S. chinensis* primarily to direct anthropogenic activity, and *N. phocaenoides* to broader water quality degradation.

A key limitation of eDNA-based approaches is that read-based relative abundance reflects a semi-quantitative signal rather than true biomass or abundance, as taxa differ in DNA shedding, degradation, transport, and amplification efficiency (Deiner et al., 2017; Kelly et al., 2014; Shelton et al., 2019). In dynamic coastal systems, hydrodynamic transport and mixing can further decouple eDNA signals from local organismal presence, introducing both false positives and false negatives. In addition, the compositional nature of read data may introduce biases when interpreting relative abundance patterns. This limitation may contribute to the weak predictive performance of our analytical approach and supports cautious interpretation of relative abundance-based inferences. Although our matched sampling design and environmental modeling reduce some of this uncertainty. Future work should explicitly validate eDNA-derived relative abundance against independent survey data and incorporate hydrodynamic context to better resolve local habitat use.

### Conservation and global relevance

Effective conservation in urbanized seascapes requires moving beyond biodiversity detection towards identifying species-specific threats and priority habitats and mechanisms of vulnerability (O’Hara et al., 2021; Sala et al., 2021; Turner et al., 2024). Integrating eDNA metabarcoding with JSDMs, our study provided a framework to distinguish environmentally structured occurrence patterns from residual predator-prey associations, highlighted divergent vulnerabilities between two co-occurring cetaceans. Unlike conventional single-species models, multispecies approaches capture shared responses and co-occurrence patterns, offering a scalable diagnostic tool in data-limited seascapes (Ovaskainen et al., 2017; Pollock et al., 2014).

Based on eDNA-JSDM findings, adaptive management actions should be tailored to prevailing pressures rather than applied uniformly. For *S. chinensis*, whose occurrence is more tightly linked to water quality and vessel activity than to prey, priority actions include stricter regulations of nutrient and contaminant inputs, turbidity controls associated with coastal developments, and improved effluent standards (Jefferson et al., 2023). *N. phocaenoides* shows strong sensitivity to elevated E. coli and nitrate nitrogen, which drive eutrophication and hypoxia and are indicative of degraded water quality and habitat suitability (Banerjee et al., 2023). Management strategies for *N. phocaenoides* should therefore prioritize improving water quality through stricter effluent controls, nutrient reduction at the land–sea interface, and monitoring microbial indicators alongside chemical pollutants. For both cetacean species, mitigating vessel disturbance through speed limits, dynamic routing, and strategic rezoning of marine protected areas remains essential (Erbe et al., 2019; Pirotta et al., 2015).

Conservation outcomes depend on tailoring actions to prevailing pressures rather than applying uniform measures. Recognizing that “one size does not fit all,” species-specific diagnostics are essential for effective management (Y. Zhao et al., 2025). Such contrasts echo global cases where uniform measures have failed. For instance, efforts for woodland caribou recovery showed that initially habitat protection plans alone were insufficient, with successful recovery efforts requiring locally tailored combinations of predator control, maternal penning, prey reduction, and habitat restoration (Serrouya et al., 2019). Other efforts to establish reef-focused MPAs have overlooked the ranges of mobile marine megafauna resulting in many species not adequately protected leading to continued population declines (Game et al., 2009; Gilmour et al., 2022; Maxwell et al., 2013). In contrast, successfully targeted interventions such as rerouting shipping lanes, implementing vessel speed restrictions that lowered lethal strikes have had positive effects on North Atlantic right whales (Nisi et al., 2024; Vanderlaan & Taggart, 2009). While regulations on gear modifications in longline fisheries have directly reduced sea turtle and albatross bycatch (Lewison et al., 2014). Our approach provides similar threat-specific diagnostics, linking biodiversity patterns directly to prevailing stressors and guiding adaptive, species-specific management. As efforts toward the Kunming–Montreal Global Biodiversity Framework’s 30-by-30 target accelerate, such approaches will be indispensable for closing conservation gaps and prioritizing effective interventions (Sala et al., 2021; Turner et al., 2024).

## Conclusion

This study demonstrates how integrating eDNA metabarcoding with multispecies distribution models can move marine conservation from monitoring to spatially targeted action. By explicitly accounting for abiotic stressors, biotic associations, and species-specific responses, our framework provides fine-scale diagnostics that reveal conservation gaps and inform adaptive management. For Hong Kong’s resident cetaceans, this means identifying refuges for *S. chinensis* that are highly sensitive to vessel traffic, and high turbidity murky water, and identifying the dependence of *N. phocaenoides* on areas with higher water quality and reduced vessel activities. These insights allow management plans to be tailored to the dominant pressures faced by each species, rather than relying on uniform conservation measures. The capacity of the eDNA-JDSM approach to identify unprotected core habitats, help disentangle overlapping environmental stressors and species responses, and priorities interventions is particularly valuable in data-limited, disturbance-prone systems where conventional monitoring is infeasible. Beyond Hong Kong, this integrative approach offers a scalable tool to advance the CBD 2030 targets by linking biodiversity detection directly to species-specific, adaptive conservation strategies.

## Acknowledgement

This project was funded by the Marine Conservation Enhancement Fund, (grant number MCEF21003 to MS). We also thank the B. Lin and R. Cheng for assistance with sampling.

## Data availability

Data available on request from the authors

