## Supplemental Material for "Pollution and Anthropogenic Stressors Are Associate with Cetacean Vulnerability in Coastal Waters: Fine-Scale Diagnostics from eDNA and Multispecies Modeling"

### Appendix

#### Appendix S1

*Full cetacean amplicon assessment: Sampling depth does not influence cetacean richness or relative abundance*

We assessed whether sampling depth or season influenced cetacean richness or relative abundance using zero-inflated Generalized Linear Mixed Models (GLMM) with a negative binomial error distribution using the glmmTMB function (glmmTMB package; [1]). Sampling ecoregions around Lantau Island were included as a random effect, with cetacean species richness as the response variable. For the relative sequence abundance of resident species, GLMMs with a Tweedie distribution were fitted using glmmTMB, including sampling site as a random intercept. For *Neophocaena phocaenoides*, model diagnostics indicated overdispersion; therefore, an additional observation-level random effect was included to account for extra-Poisson variation. Estimated marginal means and pairwise contrasts were computed using the emmeans package to assess spatiotemporal differences [2].

For the GLMMs fitted with a Tweedie error distribution, the Tweedie index parameter ($p$) was estimated by maximum likelihood and verified using profile likelihood. All estimates fell between 1 and 2, confirming that the compound Poisson–Gamma distribution was appropriate for the data. Model overdispersion was evaluated using Pearson residuals. For each model, a Pearson chi-square statistic was calculated from the sum of squared Pearson residuals and divided by the residual degrees of freedom to obtain a dispersion ratio. .

The Tweedie index parameters estimated for the spatial and seasonal models were 1.337 for *S. chinensis* and 1.477 for *N. phocaenoides*. For models evaluating environmental drivers, the index parameters were 1.404 for S. chinensis and 1.674 for *N. phocaenoides*. All values were within the range, confirming that the Tweedie distribution was appropriate for modelling zero-inflated continuous relative abundance in both species.

**Supplementary Table S1.** Environmental parameters, group variables, and their sources used in this study.

| Group (n) | Parameter (unit) | Source |
|---|---|---|
| Chemical property (15) | Biochemical Oxygen Demand (mg/L) | Current study |
| | Ammoniacal Nitrogen (mg/L) | |
| | Dissolved Oxygen (% saturation) | |
| | Dissolved Oxygen (mg/L) | |
| | Unionized Ammonia (mg/L) | |
| | Total Phosphorus (mg/L) | |
| | Total Nitrogen (mg/L) | |
| | Total Kjeldahl Nitrogen (mg/L) | |
| | Total Inorganic Nitrogen (mg/L) | |
| | Salinity (Practical Salinity Units, PSU) | |
| | Silica (mg/L) | |
| | Acidity/Alkalinity (pH) | |
| | Orthophosphate Phosphorus (mg/L) | |
| | Nitrite-Nitrogen (mg/L) | |
| | Nitrate-Nitrogen (mg/L) | |
| Physical property (5) | Volatile Suspended Solids (mg/L) | |
| | Turbidity (Nephelometric Turbidity Units) | |
| | Water Temperature (°C) | |
| | Suspended Solids (mg/L) | |
| | Secchi Disc Depth (m) | |
| Biological property (4) | Phaeopigments (g/L) | |
| | Faecal Coliforms (colony-forming units per 100 mL) | |
| | Escherichia coli (colony-forming units per 100 mL) | |
| | Chlorophyll-a (µg/L) | |
| Anthropogenic disturbance | Vessel presence (hours/0.5 $km^2$) | Global Fishing Watch |
| | Apparent fishing effort (hours/0.5 $km^2$) | |

**Supplementary Table S2.** Vessel presence and apparent fishing effort across ecoregions during the rainy and dry seasons.

| ID | Ecoregion | Site | Depth | Season | Vessel presence | Apparent fishing effort |
|---|---|---|---|---|---|---|
| MCEF_1_1_B_X2 | NLI | 1 | Bottom | Dry season | 471 | 4.99 |
| MCEF_1_1_M_X2 | NLI | 1 | Mid | Dry season | 471 | 4.99 |
| MCEF_1_1_S_X2 | NLI | 1 | Surface | Dry season | 471 | 4.99 |
| MCEF_1_2_M_X2 | NLI | 2 | Mid | Dry season | 46 | 0 |
| MCEF_1_2_S_X2 | NLI | 2 | Surface | Dry season | 46 | 0 |
| MCEF_1_3_B_X2 | NLI | 3 | Bottom | Dry season | 18 | 0 |
| MCEF_1_3_M_X2 | NLI | 3 | Mid | Dry season | 18 | 0 |
| MCEF_1_3_S_X2 | NLI | 3 | Surface | Dry season | 18 | 0 |
| MCEF_1_4_M_X2 | NLI | 4 | Mid | Dry season | 544 | 1.62 |
| MCEF_1_4_S_X2 | NLI | 4 | Surface | Dry season | 544 | 1.62 |
| MCEF_2_1_B_X2 | ELI | 1 | Bottom | Dry season | 843 | 0.48 |
| MCEF_2_1_M_X2 | ELI | 1 | Mid | Dry season | 843 | 0.48 |
| MCEF_2_1_S_X2 | ELI | 1 | Surface | Dry season | 843 | 0.48 |
| MCEF_2_2_M_X2 | ELI | 2 | Mid | Dry season | 948 | 0 |
| MCEF_2_2_S_X2 | ELI | 2 | Surface | Dry season | 948 | 0 |
| MCEF_2_3_B_X2 | ELI | 3 | Bottom | Dry season | 18 | 0 |
| MCEF_2_3_M_X2 | ELI | 3 | Mid | Dry season | 18 | 0 |
| MCEF_2_3_S_X2 | ELI | 3 | Surface | Dry season | 18 | 0 |
| MCEF_2_4_M_X2 | ELI | 4 | Mid | Dry season | 133 | 1.12 |
| MCEF_2_4_S_X2 | ELI | 4 | Surface | Dry season | 133 | 1.12 |
| MCEF_3_1_B_X2 | SELI | 1 | Bottom | Dry season | 3 | 0 |
| MCEF_3_1_M_X2 | SELI | 1 | Mid | Dry season | 3 | 0 |
| MCEF_3_1_S_X2 | SELI | 1 | Surface | Dry season | 3 | 0 |
| MCEF_3_2_B_X2 | SELI | 2 | Bottom | Dry season | 1 | 0 |
| MCEF_3_2_M_X2 | SELI | 2 | Mid | Dry season | 1 | 0 |
| MCEF_3_2_S_X2 | SELI | 2 | Surface | Dry season | 1 | 0 |
| MCEF_3_3_M_X2 | SELI | 3 | Mid | Dry season | 6 | 1.01 |
| MCEF_3_4_B_X2 | SELI | 4 | Bottom | Dry season | 6 | 1.01 |
| MCEF_3_4_M_X2 | SELI | 4 | Mid | Dry season | 6 | 1.01 |
| MCEF_3_4_S_X2 | SELI | 4 | Surface | Dry season | 6 | 1.01 |
| MCEF_4_1_M_X2 | SLI | 1 | Mid | Dry season | 6 | 0 |
| MCEF_4_2_B_X2 | SLI | 2 | Bottom | Dry season | 1 | 0 |
| MCEF_4_2_M_X2 | SLI | 2 | Mid | Dry season | 1 | 0 |
| MCEF_4_2_S_X2 | SLI | 2 | Surface | Dry season | 1 | 0 |
| MCEF_4_3_M_X2 | SLI | 3 | Mid | Dry season | 1 | 0 |

| | | | | | | |
|---|---|---|---|---|---|---|
| MCEF_4_4_B_X2 | SLI | 4 | Bottom | Dry season | 2 | 2 |
| MCEF_4_4_M_X2 | SLI | 4 | Mid | Dry season | 2 | 2 |
| MCEF_4_4_S_X2 | SLI | 4 | Surface | Dry season | 2 | 2 |
| MCEF_5_1_M_X2 | SWLI | 1 | Mid | Dry season | 5 | 0 |
| MCEF_5_2_B_X2 | SWLI | 2 | Bottom | Dry season | 1 | 0 |
| MCEF_5_2_M_X2 | SWLI | 2 | Mid | Dry season | 1 | 0 |
| MCEF_5_2_S_X2 | SWLI | 2 | Surface | Dry season | 1 | 0 |
| MCEF_5_3_M_X2 | SWLI | 3 | Mid | Dry season | 1 | 0 |
| MCEF_5_4_M_X2 | SWLI | 4 | Mid | Dry season | 4 | 0 |
| MCEF_1_1_M_X1 | NLI | 1 | Mid | Rainy season | 57 | 0 |
| MCEF_1_2_M_X1 | NLI | 2 | Mid | Rainy season | 38 | 0 |
| MCEF_1_3_M_X1 | NLI | 3 | Mid | Rainy season | 0 | 0 |
| MCEF_1_4_M_X1 | NLI | 4 | Mid | Rainy season | 179 | 1.12 |
| MCEF_2_1_B_X1 | ELI | 1 | Bottom | Rainy season | 472 | 0.79 |
| MCEF_2_1_M_X1 | ELI | 1 | Mid | Rainy season | 472 | 0.79 |
| MCEF_2_1_S_X1 | ELI | 1 | Surface | Rainy season | 472 | 0.79 |
| MCEF_2_2_M_X1 | ELI | 2 | Mid | Rainy season | 1304 | 0 |
| MCEF_2_3_M_X1 | ELI | 3 | Mid | Rainy season | 34 | 0 |
| MCEF_2_4_M_X1 | ELI | 4 | Mid | Rainy season | 629 | 0 |
| MCEF_3_1_B_X1 | SELI | 1 | Bottom | Rainy season | 22 | 0 |
| MCEF_3_1_M_X1 | SELI | 1 | Mid | Rainy season | 22 | 0 |
| MCEF_3_1_S_X1 | SELI | 1 | Surface | Rainy season | 22 | 0 |
| MCEF_3_2_M_X1 | SELI | 2 | Mid | Rainy season | 26 | 4.46 |
| MCEF_3_2_S_X1 | SELI | 2 | Surface | Rainy season | 26 | 4.46 |
| MCEF_3_3_M_X1 | SELI | 3 | Mid | Rainy season | 13 | 4.67 |
| MCEF_3_4_M_X1 | SELI | 4 | Mid | Rainy season | 102 | 11.37 |
| MCEF_4_1_B_X1 | SLI | 1 | Bottom | Rainy season | 8 | 0 |
| MCEF_4_1_M_X1 | SLI | 1 | Mid | Rainy season | 8 | 0 |
| MCEF_4_1_S_X1 | SLI | 1 | Surface | Rainy season | 8 | 0 |
| MCEF_4_2_B_X1 | SLI | 2 | Bottom | Rainy season | 8 | 0 |
| MCEF_4_2_M_X1 | SLI | 2 | Mid | Rainy season | 8 | 0 |
| MCEF_4_2_S_X1 | SLI | 2 | Surface | Rainy season | 8 | 0 |
| MCEF_4_3_B_X1 | SLI | 3 | Bottom | Rainy season | 5 | 0.56 |
| MCEF_4_3_M_X1 | SLI | 3 | Mid | Rainy season | 5 | 0.56 |
| MCEF_4_3_S_X1 | SLI | 3 | Surface | Rainy season | 5 | 0.56 |
| MCEF_4_4_M_X1 | SLI | 4 | Mid | Rainy season | 2 | 2.78 |
| MCEF_5_1_B_X1 | SWLI | 1 | Bottom | Rainy season | 1 | 0.27 |
| MCEF_5_1_M_X1 | SWLI | 1 | Mid | Rainy season | 1 | 0.27 |
| MCEF_5_1_S_X1 | SWLI | 1 | Surface | Rainy season | 1 | 0.27 |
| MCEF_5_2_M_X1 | SWLI | 2 | Mid | Rainy season | 3 | 0 |
| MCEF_5_3_M_X1 | SWLI | 3 | Mid | Rainy season | 4 | 0 |
| MCEF_5_4_M_X1 | SWLI | 4 | Mid | Rainy season | 4 | 0 |

**Supplementary Table S3.** The summary statistics of post-hoc pairwise PERMANOVA, comparing the differences the season and the dwelling depth of cetacean communities.

| **Comparison** | ***F*-value** | **$R^2$** | **Adjusted *p*-value** | **Significance** |
|---|---|---|---|---|
| Dry season mid-water vs Rainy season mid-water | 1.89 | 0.07 | 0.117 | ns |
| Dry season mid-water vs Rainy season surface | 0.68 | 0.03 | 0.64 | ns |
| Dry season mid-water vs Rainy season bottom | 2.14 | 0.13 | 0.059 | ns |
| Dry season mid-water vs Dry season bottom | 0.72 | 0.07 | 0.681 | ns |
| Dry season mid-water vs Dry season surface | 0.55 | 0.06 | 0.812 | ns |
| Rainy season mid-water vs Rainy season surface | 0.69 | 0.02 | 0.605 | ns |
| Rainy season mid-water vs Rainy season bottom | 0.59 | 0.02 | 0.657 | ns |
| Rainy season mid-water vs Dry season bottom | 2.67 | 0.12 | 0.027 | • |
| Rainy season mid-water vs Dry season surface | 2.1 | 0.1 | 0.09 | ns |
| Rainy season surface vs Rainy season bottom | 0.89 | 0.04 | 0.48 | ns |
| Rainy season surface vs Dry season bottom | 1.36 | 0.09 | 0.195 | ns |
| Rainy season surface vs Dry season surface | 1.22 | 0.08 | 0.292 | ns |
| Rainy season bottom vs Dry season bottom | 2.58 | 0.22 | 0.016 | • |
| Rainy season bottom vs Dry season shallow | 2.47 | 0.22 | 0.022 | • |
| Dry season bottom vs Dry season surface | 0.19 | 0.05 | 0.9 | ns |

**Supplementary Table S4.** The summary statistics of post-hoc pairwise PERMANOVA, comparing the differences the season and the dwelling depth of fish communities.

| **Comparison** | ***F*-value** | **$R^2$** | **Adjusted** | **Significance** |
|---|---|---|---|---|

| | | | *p*-value | |
|---|---|---|---|---|
| Dry season mid-water vs Rainy season mid-water | 13.57 | 0.26 | 0.001 | ** |
| Dry season mid-water vs Rainy season surface | 8.91 | 0.23 | 0.001 | ** |
| Dry season mid-water vs Rainy season bottom | 7.93 | 0.23 | 0.001 | ** |
| Dry season mid-water vs Dry season bottom | 1.59 | 0.06 | 0.11 | ns |
| Dry season mid-water vs Dry season surface | 0.79 | 0.03 | 0.641 | ns |
| Rainy season mid-water vs Rainy season surface | 0.64 | 0.02 | 0.778 | ns |
| Rainy season mid-water vs Rainy season bottom | 0.96 | 0.04 | 0.455 | ns |
| Rainy season mid-water vs Dry season bottom | 10.86 | 0.31 | 0.001 | ** |
| Rainy season mid-water vs Dry season surface | 8.85 | 0.26 | 0.001 | ** |
| Rainy season surface vs Rainy season bottom | 0.69 | 0.04 | 0.727 | ns |
| Rainy season surface vs Dry season bottom | 8.33 | 0.34 | 0.002 | * |
| Rainy season surface vs Dry season surface | 6.15 | 0.27 | 0.001 | ** |
| Rainy season bottom vs Dry season bottom | 8.38 | 0.41 | 0.001 | ** |
| Rainy season bottom vs Dry season surface | 5.76 | 0.31 | 0.001 | ** |
| Dry season bottom vs Dry season surface | 1.19 | 0.10 | 0.284 | ns |

### Supplementary Figures

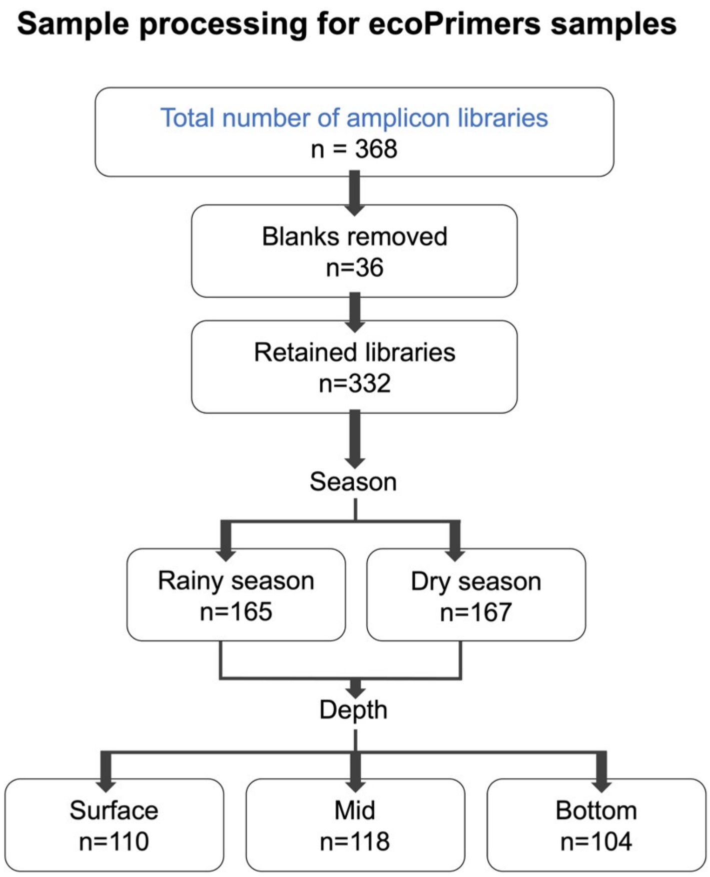


**Supplementary Figure S1.** Workflow for sample selection and quality control of the full (left) and subset (right) seawater eDNA datasets targeting cetaceans using the ecoPrimers set.

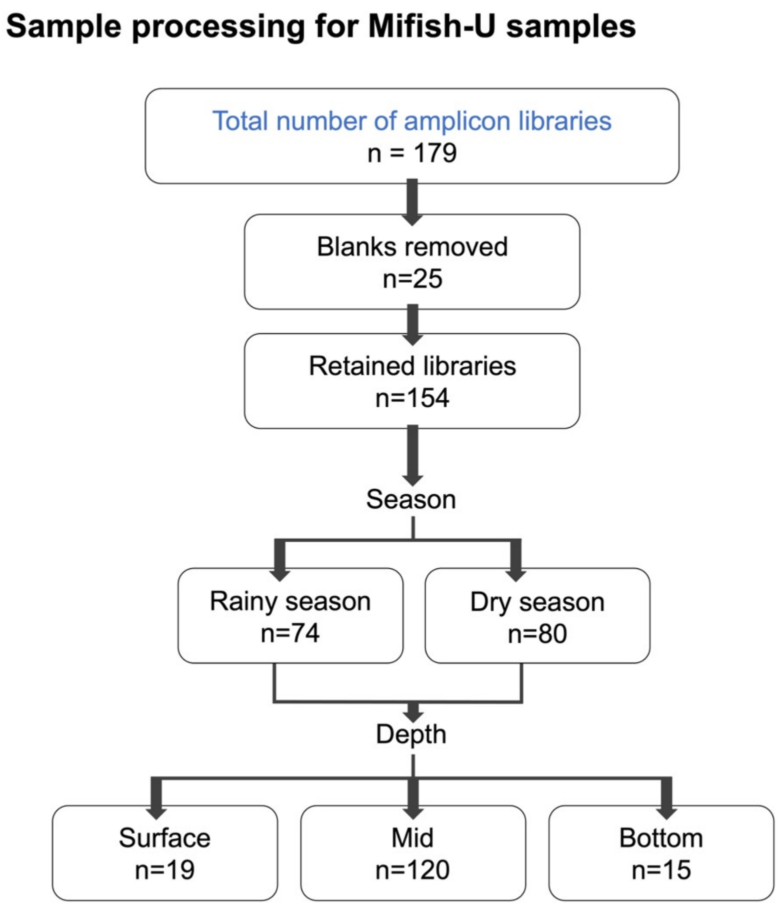


**Supplementary Figure S2.** Workflow for sample selection and quality control of the seawater eDNA datasets targeting fish using the MiFish-U primers set.

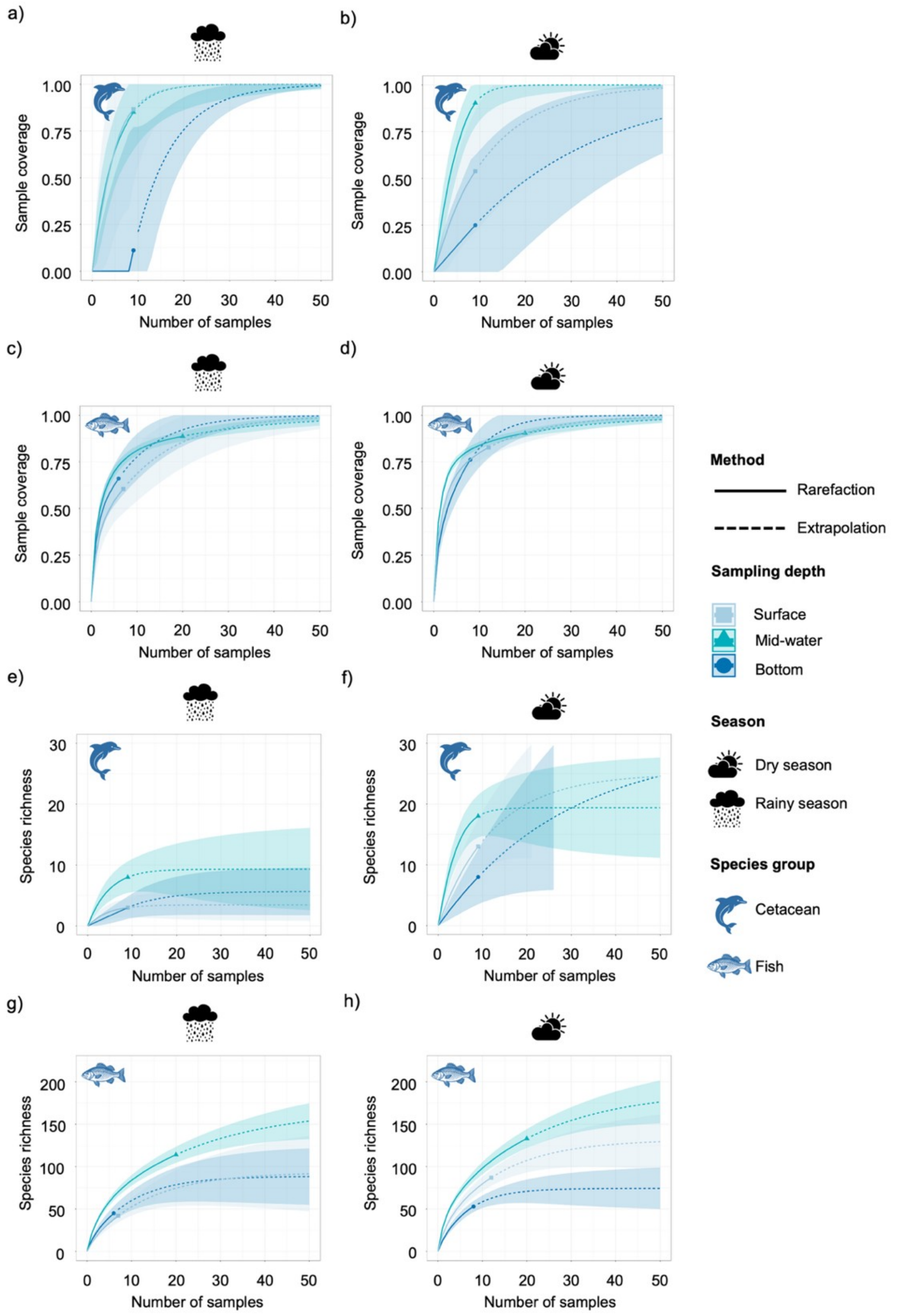


**Supplementary Figure S3.** Rarefaction and extrapolation curves for sampling coverage (Cetaceans: a, b; Fish: c, d) and species richness (Cetaceans: e, f; Fish: g, h) using Hill number q = 0. Results are presented by season (left column: *rainy*; right column: *dry*) and depth. Solid lines represent observed values; dashed lines represent extrapolated values. Shaded areas and dotted lines indicate 95% confidence intervals.

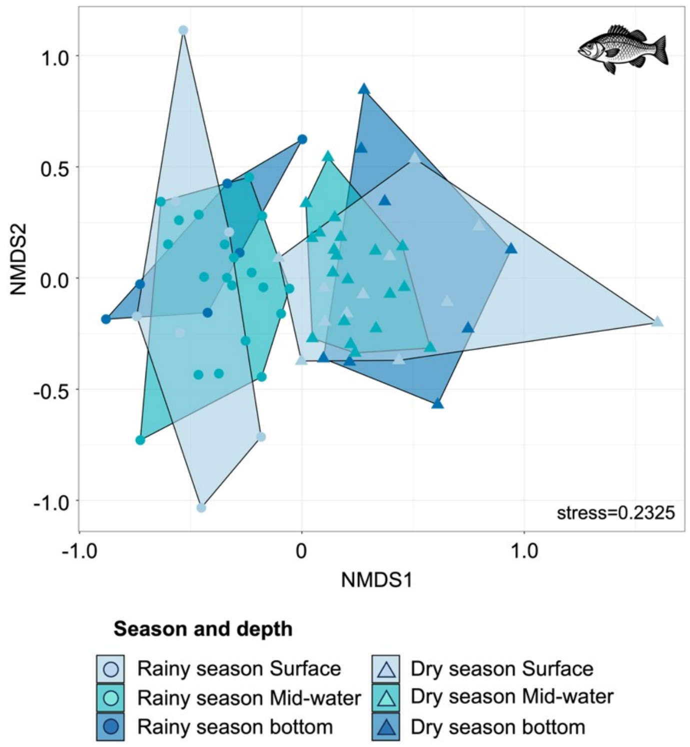


**Supplementary Figure S4.** Nonmetric Multidimensional Scaling (NMDS) ordination generated using the Sørensen dissimilarity measure based on the occurrence of fish.

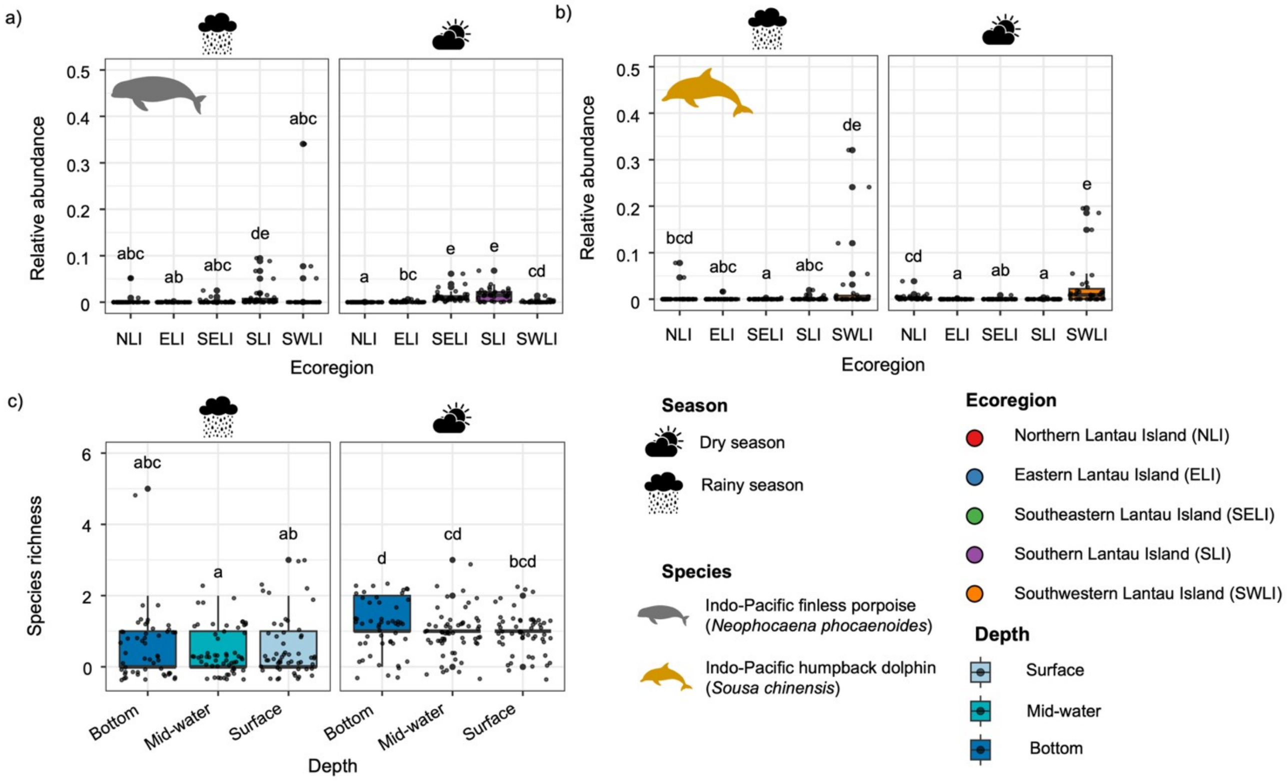


**Supplementary Figure S5.** Spatiotemporal variation in the relative abundance of resident cetaceans and total cetacean species richness from the larger dataset. Panels show the relative abundance of (a) Indo-Pacific finless porpoises (*Neophocaena phocaenoides*) and (b) Indo-Pacific humpback dolphins (*Sousa chinensis*) during the rainy and dry seasons across sampling locations. Panel (c) presents total cetacean species richness across sampling strata and seasons. Compact letters indicate significant spatiotemporal differences in relative abundance ($p < 0.05$).

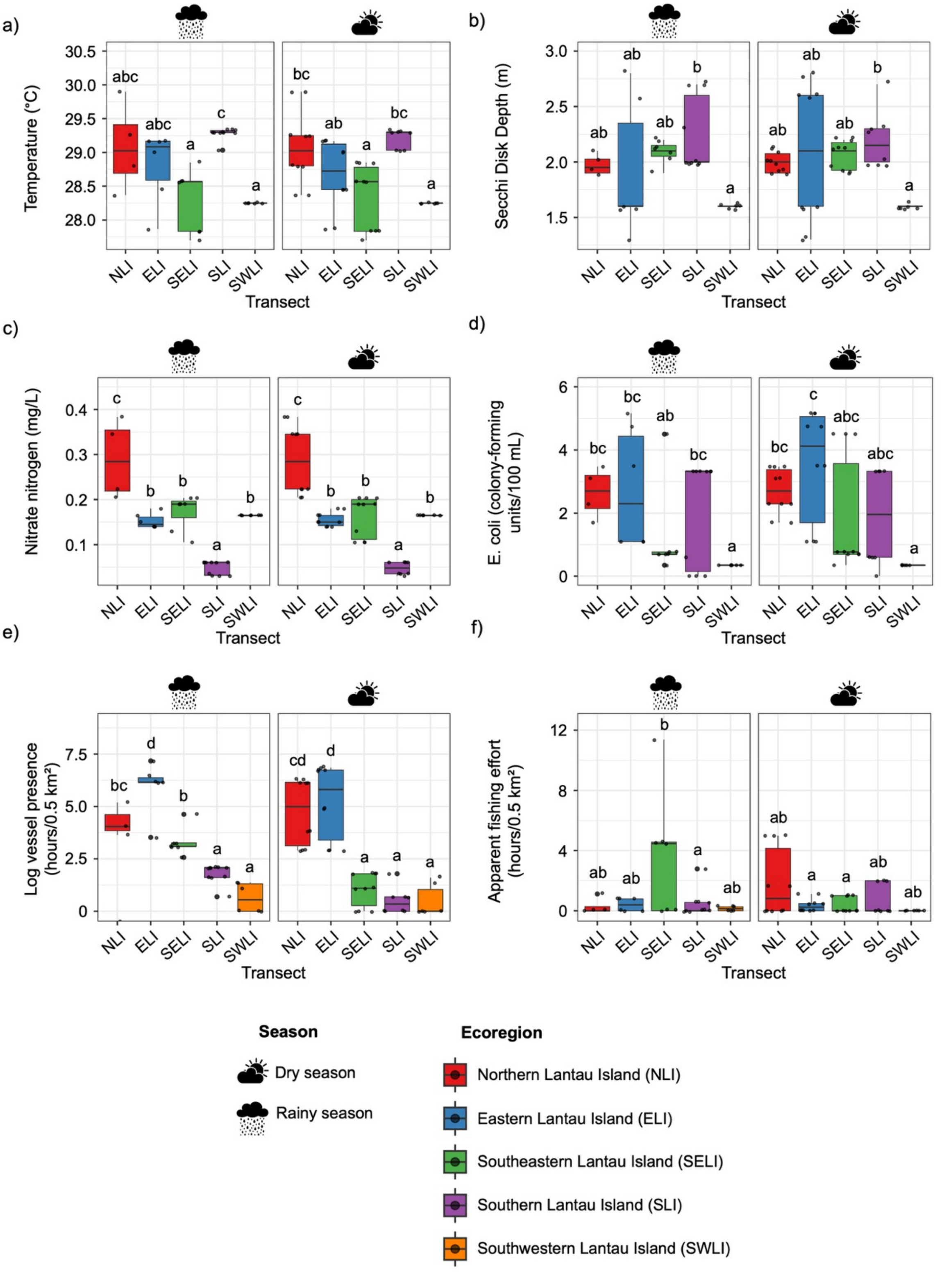


**Supplementary Figure *S*6.** *Spatiotemporal* variation in filtered environmental covariates across ecoregions. Compact letters indicate statistically significant differences among groups ($p < 0.05$), based on pairwise comparisons.

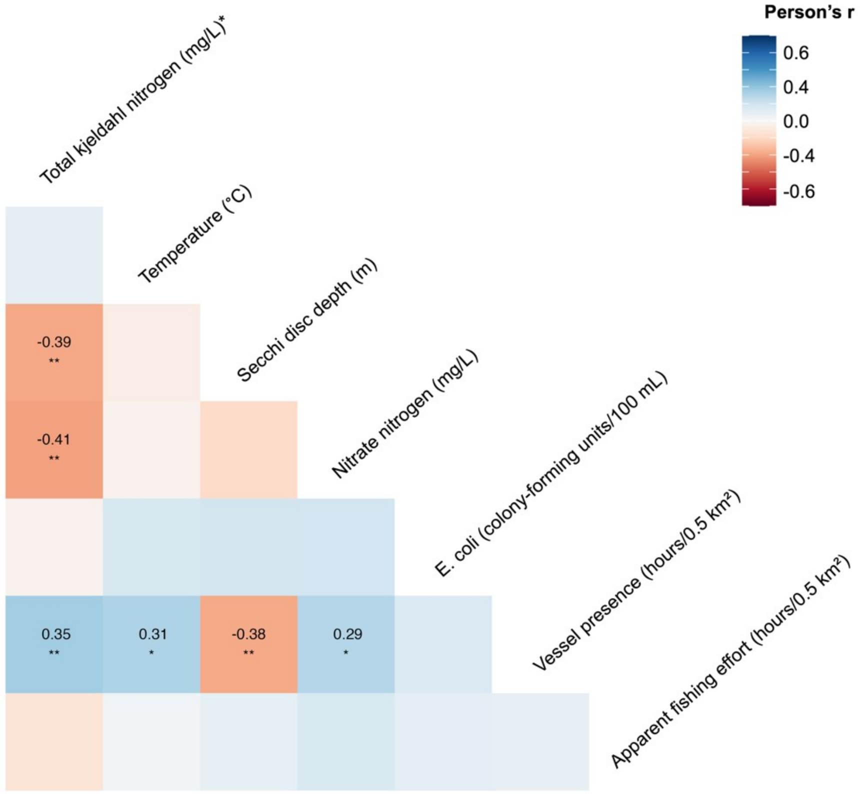


**Supplementary Figure S7.** Correlation matrix of environmental covariates after filtering. Environmental variables were filtered using pairwise correlation ($|r| \geq 0.7$) and variance inflation factor (VIF $\geq 5$) thresholds to reduce multicollinearity. Variables marked with an asterisk (*) were excluded from subsequent analyses to improved HMSC model performance.

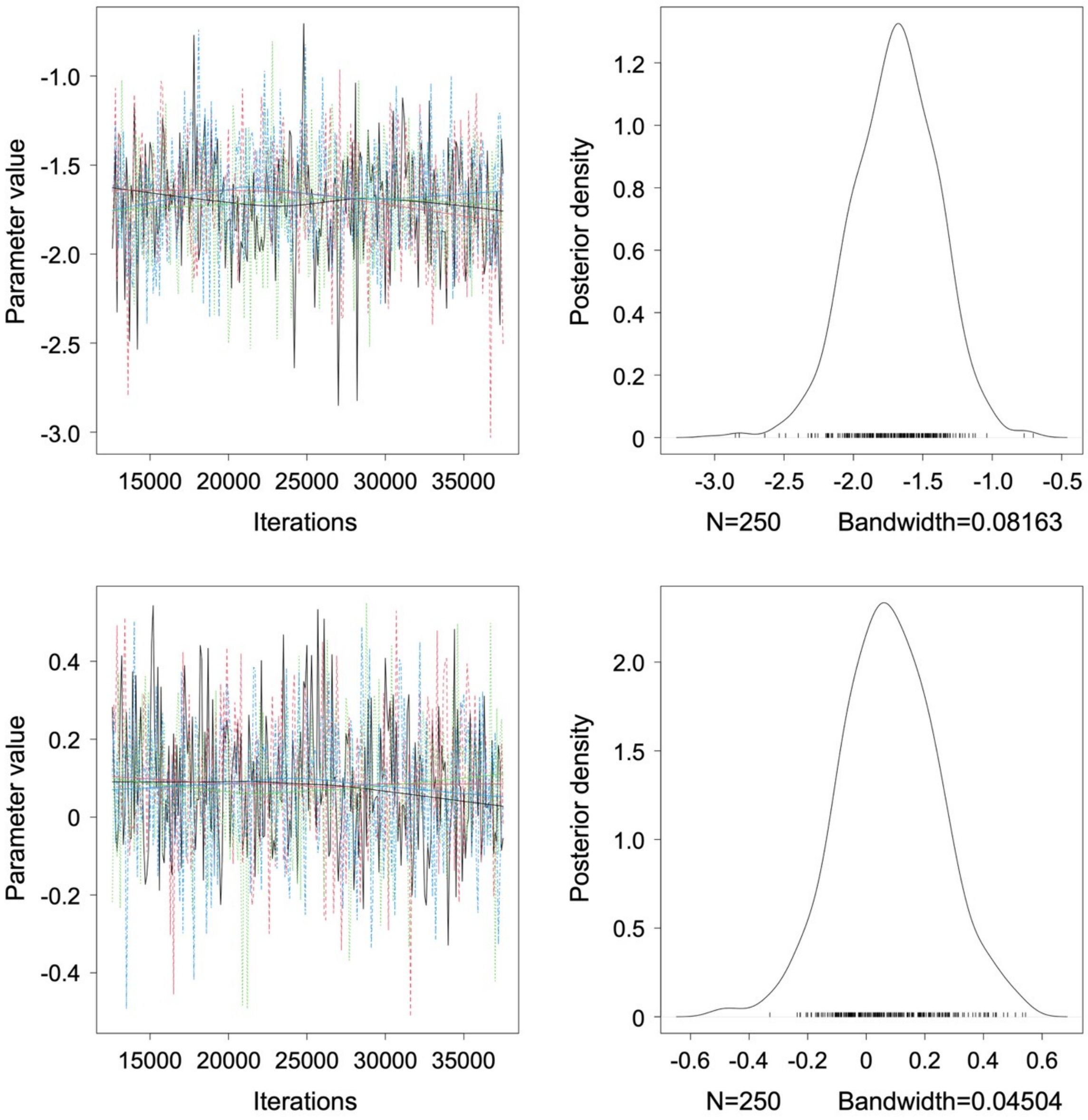


**Supplementary Figure S8.** Posterior trace plots and density estimate for the $\beta$ parameters. The left column shows the MCMC trace plots for the $\beta$ parameters, and the right column shows the corresponding posterior density estimates. The top panels present results from the null model (intercept-only), while the bottom panels show results from the full model, which includes environmental covariates.

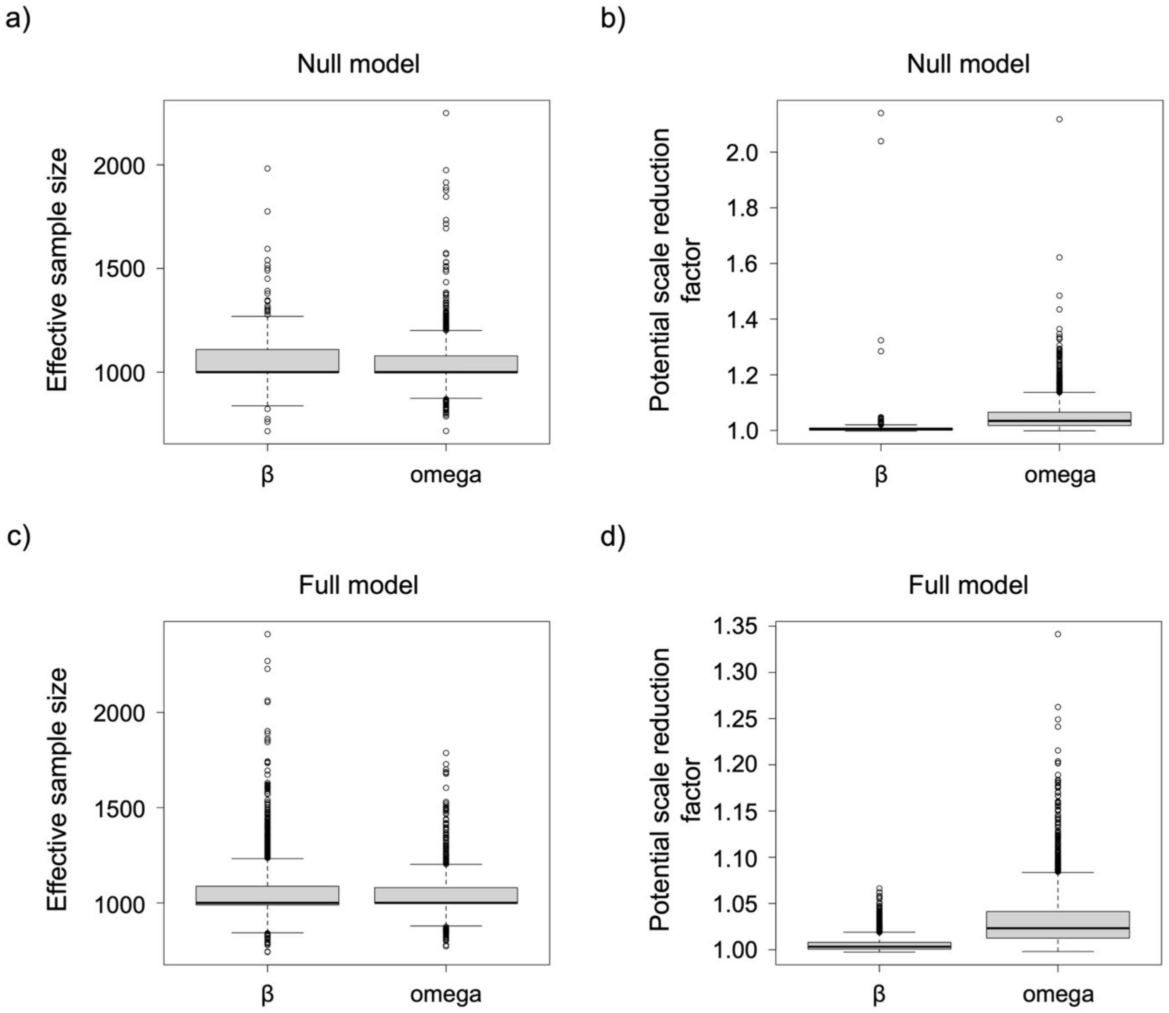


**Supplementary Figure S9.** Diagnostic boxplots for MCMC convergence of the HMSC models, showing results for both the null and full models. Panels (a, c) display the effective sample size, while panels (b, d) show the potential scale reduction factor (PSRF). Each plot summarizes convergence for the $\beta$ parameters (species responses to environmental covariates) and the omega parameters (predator –prey associations at the ecoregion level).

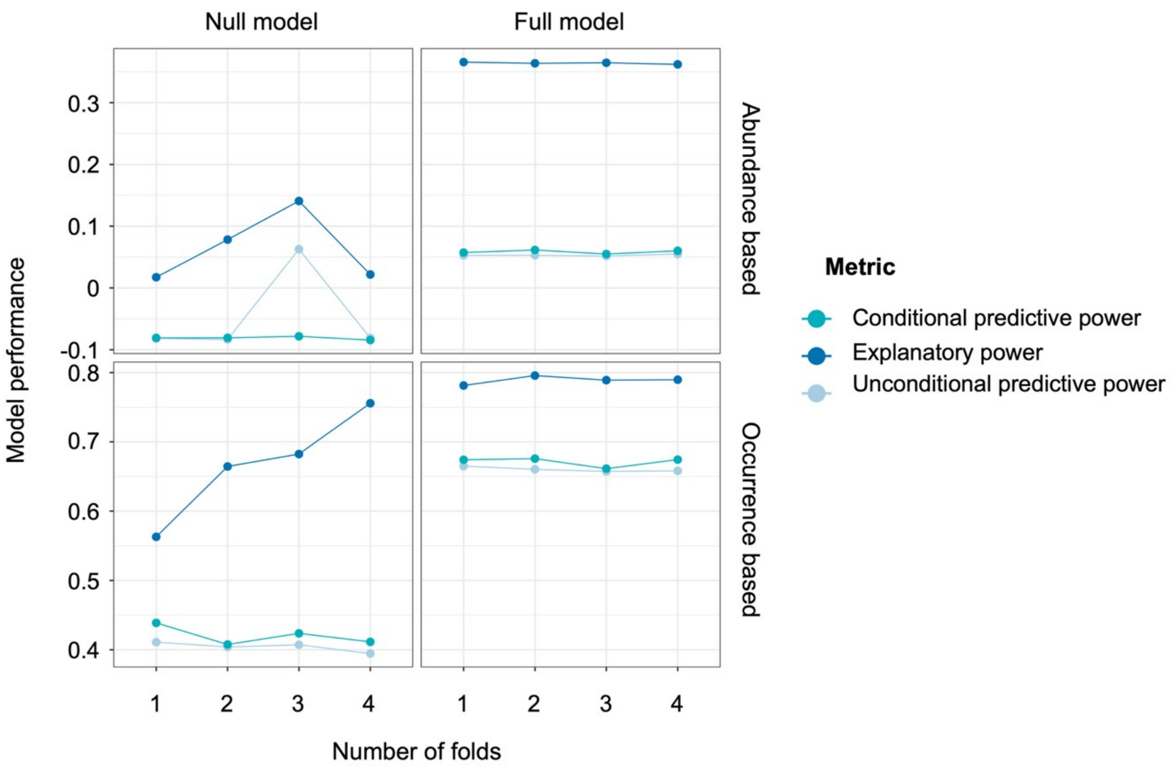


**Supplementary Figure S10.** Sensitivity of model performance to cross-validation design. Explanatory, unconditional predictive, and conditional predictive performance are shown across increasing numbers of cross-validation folds (nf) for null and full models (columns) and for abundance and presence–absence responses (rows). Points and lines indicate mean performance across species. Predictive and conditional metrics vary little with nf, indicating that model inference is robust to the choice of cross-validation fold number.

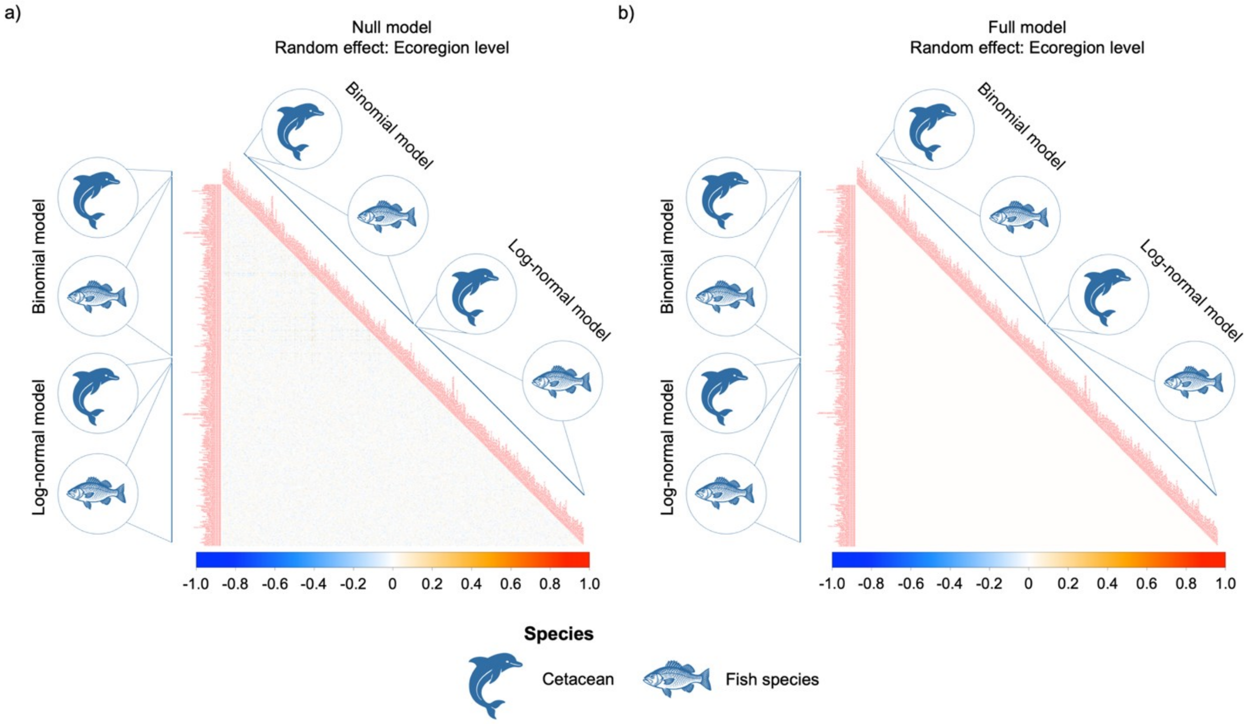


**Supplementary Figure S11.** Species–species association matrix estimated from the HMSC models. Panel (a) shows associations from the null model, representing raw predator-prey associations, and (b) shows associations from the full model, constrained with environmental predictors at the ecoregion random level. Each matrix includes derived from both binomial (occurrence) and log-normal (abundance) models. Only associations with posterior support ≥ 0.75 (positive) or ≤ 0.25 (negative) are shown. The colour gradient represents both the direction and magnitude of associations, with deeper hues reflecting stronger interspecific relationships.
